# Correlated Evolution of Metabolic Functions over the Tree of Life

**DOI:** 10.1101/093591

**Authors:** Murray Patterson, Thomas Bernard, Daniel Kahn

## Abstract

We are interested in the structure and evolution of metabolism in order to better understand its complexity. We study metabolic functions in 1459 species within which several hundreds of thousands of families of homologous genes have been identified [17]. Given a protein sequence, PRIAM search [5] delivers probabilities of the presence of several thousand enzymes (ECs). This allows us to infer reaction sets and to construct a metabolic network for an organism, given its set of sequences.

We then propagate these ECs to the ancestral nodes of the species tree using maximimum likelihood methods. These evolutionary scenarios are systematically compared using pairwise mutual information. We identify co-evolving enzyme sets from the graph of these relationships using community detection algorithms [1,3]. This sheds light on the structure of the metabolic networks in terms of co-evolving metabolic modules. These modules are also interpreted from a functional perspective using stoichiometric models of metabolic networks.

## Introduction

The functioning of a cell or an organism is a combination of reactions operating on compounds that together form a complex network. At any given level of detail in this network, one can find functional sub-units – subsets of highly interconnected reactions – from basic processes involving just a handful of reactions *e.g.,* the Calvin cycle in photosynthesis, all the way to major building blocks of a cell, like a ribosome. The work of Ravasz *et al.* [18] is an important large-scale study showing that metabolic networks are indeed hierarchically organized sets of such sub-units, or *modules.* Today, this notion is well-known, and we have established standards such as the KEGG resource [11].

These modules, and their organization, are important indicators of the evolutionary processes that the cell/organism has undergone throughout time. For example, one would see a high degree of selection throughout time for important basic modules like the Calvin cycle, while the event where plants first acquired the chloroplast from cyanobacteria [15] would be marked by the emergence of this module in the plant clade. Following the work of [18], large-scale studies emerged that are similar in spirit to [18], but focusing on modularity from an evolutionary point of view. The first such study was not long after the release of the KEGG resource, where some of its authors extracted phylogenetically related sub-units from the modules in KEGG [20,21], which they refer to as *phylogenetic network modules.* A few years later, Kim and Price [12] take a similar approach where they try to find phylogenetically related genes based on co-occurrence patterns in the genomes of a set of microbes. In this work, we perform a study that is more similar to [21], the big difference being that our approach is *unsupervised*: we simply extract phylogenetically related sub-units in general, and not in the context of currently existing modules. There are, of course, other important differences to [21] and [12] of our approach, that we note as we detail this work in the following.

Our study is on a set of 1463 organisms from the tree of life, including bacteria, archaea and eukaryotes, for which there is a species phylogeny (see Supplementary Figure 1 for a graphical representation of this phylogeny). These organisms (genome sequences) are taken from the HOGENOM database [17], *i.e.,* they are composed of several hundreds of thousands of families of homologous genes. Similarily to [21], we reason about the presence/absence of a metabolic reaction in terms of the *enzymes* that catalyze said reaction. We infer the presence/absence of enzymes in an organism based on the *protein sequence* content encoded by its genome. We do this using PRIAM [5] – a set of *profiles,* or *position-specific scoring matrices* (PSSMs) for protein modules covering the Swiss-Prot database. Combining the results of the rps-blast(s) of the profile(s) for a given enzyme, or *enzyme code* (EC), against all protein sequences encoded by a genome delivers a probability that the given EC is present in the organism. So, in fact, we have a probabilistic framework, which is more general than just the (binary) presence/absence data, as used by both [21] and [12].

In addition to using binary data, [21] and [12] compare *phylogenetic profiles* (the presence/absence data for each organism) directly to assess the correlation between a pair of ECs and genes, respectively. Due, simply to *phylogenetic inertia*: the propensity of a character state to remain the same from ancestor to decendent, a pair of ECs that both appear as present early in the phylogeny for a large clade would tend to be co-present in a large number of leaves of this clade, rendering it much more highly correlated, from a phylogenetic profile point of view, than a pair of states that both appear later in the phylogeny. In this work, we first use a maximum likelihood method [7] to propagate gain and loss probabilities of an EC to the branches of the species phylogeny, given the probabilities of this EC at the leaves. The correlation of a pair of ECs is then in terms of its gain (and loss) probabilities on the branches. Here, each correlated gain (or loss) will be weighted equally, regardless of where it occurred in the phylogeny, avoiding this bias by phylogenetic inertia.

More specifically, the correlation between a pair of ECs is computed as the *mutual information* (MI) of the gain (and loss) probabilities of each EC over the branches of the tree. The notion of mutual information, coming from Shannon information theory, is a measure of the *joint entropy* of a pair of distributions, and mutual information can be no higher than the entropy of either of the distributions alone. That is to say that if an EC (or pair of ECs) is very prevalent in the phylogeny, it will have a low entropy, and hence the MI of this EC with another (or this pair of ECs) will not be so high that one needs to resort to filtering, as they mention what is necessary in [12]. The MI between a pair of ECs is a measure of the degree to which this pair is co-evolving throughout evolutionary history. We compare MI for a pair of ECs to the degree to which this pair is co-evolving as computed by [16] and find a correlation between these two measures, validating this approach.

Finally, we want to find sets of ECs that are co-evolving, *i.e.,* these (unsupervised) phylogenetically related sub-units. To do this, we build a graph on the mutual information relationship between pairs of ECs and then find clusters of ECs in this graph using community detection algorithms [1,3]. Even though these clusters are unsupervised, *i.e.,* they are based purely on co-evolution, without any prior knowledge of how they are structured in the metabolic networks of extant organisms, we compare them to KEGG modules and find a decent degree of agreement between the two. This gives evidence that our approach has promise: that the resulting clusters contain useful information about the evolution of metabolism.

Similarily to [20,21], we reason about the presence of a metabolic reaction in an organism in terms of the *enzymes* that catalyze said reaction. We infer the presence of enzymes in an organism based on the *protein sequence* content encoded by its genome. In order to do this, *i.e.,* annotate organisms with information about function, we use Priam [5].

What we do is first annotate the extant organisms with information about function, in terms of enzyme presence probability, and then we propagate this information to the ancestral nodes of the phylogeny using maximum likelihood methods.

We try to assess the degree to which our 2732 enzymes are correlated using purely phylogenetic profiles (without taking into account the species phylogeny), since many methods take this approach [12,21].

This method is similar to our MapNH/MI pipeline, in that it takes the same inputs, namely a phylogeny and a set of states, and returns a value indicating the correlatedness of pairs of characters. We hence compare the two methods in what follows.

## Materials and Methods

### Obtaining the proteomes of the organisms

In this study, we obtain the complete proteome for the 1452 organisms listed in S1 Table. Our source of information is the Database of Complete Genome Homologous Gene Families (Hogenom) [17], release 6, found at http://pbil.univ-lyon1.fr/databases/ This release contains protein sequences for over one thousand fully sequenced organisms from eukarya, bacteria and archaea. From this release, we extract the protein sequences for all of its organisms, in the form of fasta files, and do some processing to obtain the complete proteomes for the above 1452 organisms. This procedure is detailed in S1 Text.

### Enzyme assignment with Priam

For each of the 1452 organisms listed in S1 Table, we use Priam [5] to predict the enzymes that are present in each organism, given its proteome, in fasta format.

Priam is a collection of *profiles,* or position-specific scoring matrices that were trained using the Swiss-Prot database. The idea is that a blast of the profiles against a protein sequence delivers presence probabilities of each enzyme that is predictable by Priam. In order to obtain an overall probability of presence of each enzyme in an organism, given the proteome, we use the Priam search tool. This tool performs the blasts of the profiles against each sequence in the proteome and then combines the results for each enzyme to deliver this desired overall probability in the organism. In addition to this, it can also use this information to construct a draft metabolic network, and corresponding stoichiometric matrix for this organism.

We downloaded the Feb 2014 release of Priam from `priam.prabi.fr`, and the latest search tool, and then for each of the 1452 organisms listed in S1 Table, used it to predict the probability of presence of each of the enzymes predictable by Priam. We then used Priam search to construct a draft metabolic network for each of these 1452 organisms. Because our study is more concerned with core metabolism than secondary metabolism, we do not consider enzymes that exlusively catalyze macromolecular reactions, restricting our study to the 2732 enzymes listed in S2 Table. Details on how we select these 2732 enzymes among those predictable by Priam is detailed in S2 Text. This means that, for each of these 2732 enzymes, we have a probability of its presence in each of our 1452 organisms, what we refer to as a *phylogenetic profile* of the enzyme.

### The phylogenetic tree

We then infer an ultrametric phylogenetic tree on our 1452 organisms listed in S1 Table, as depicted in S1 Fig. Details about how we infer this phylogenetic tree are detailed in S3 Text.

### Evolutionary scenarios

Given the phylogenetic profile for each of the 2732 enzymes computed in Enzyme assignment with Priam and the ultrametric phylogenetic tree on the 1452 organisms computed in The phylogenetic tree, we input this pair into MapNH [7] in order to infer the expected number of *gains* and *losses* of the enzyme on each branch of the phylogenetic tree. A graphical representation of the expected number of gains and losses on the phylogenetic tree is given in S2 Fig and S3 Fig.

Because MapNH [7] only supports discrete states, and our phylogenetic profiles consist of probabilities, we needed to modify MapNH to allow the input of probabilities. MapNH is a part of the suite of programs called TestNH which uses the framework called Bio++, both found at biopp.univ-montp2.fr/. We modified several parts of Bio++ to allow the input of probabilistic states to some of the maximum likelihood methods that use Bio++, including MapNH. One can use S1 Script to download and install this modified version with several different options.

This modification also includes the output of a probability (instead of the default: expected number) of gain(s) and loss(es) of a discrete character, such as an enzyme, on the branches of the phylogenetic tree. We compute these as well, for each of our 2732 enzymes, and refer to the resulting vector of gain (resp., loss) probabilities over the branches of the phylogenetic tree as a *gain evolutionary scenario* (resp., *loss evolutionary scenario*) of the enzyme.

Finally, BppAncestor [6] is another maximum likelihood method that uses Bio++ to infer the probabilities of character states at the ancestral nodes of a phylogenetic tree. It benefits from the aformentioned modification to Bio++, and so for each enzyme, we also input its phylogenetic profile and the phylogenetic tree into BppAncestor to infer the probabilities of the enzyme at each ancestral node of the phylogenetic tree.

### Mutual information

To asses how concerted the evolution of a pair of enzymes is, we compute the mutual information of their respective phylogenetic profiles computed in Enzyme assignment with Priam, and the mutual information of their (gain and loss) evolutionary scenarios computed in Evolutionary scenarios. Both consist of the computation of the mutual information of a pair vectors of probabilities, which we define in the following.

First, the overall probability *P*(*v*) of a vector *v* = [*p*_1_,…,*p_n_*] of probabilities is simply the normalized sum

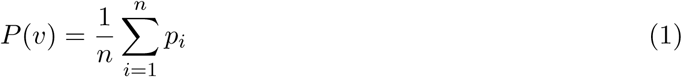

of this vector *v*. The overall joint probability *P*(*v*, *u*) of *v* with vector *u* = [*q*_1_,…, *q_n_*] of probabilities is the normalized sum of the scalar product

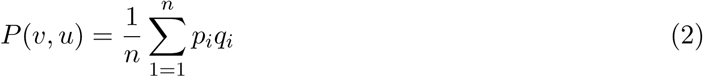

of vector *v* with *u*. Note that Equation 2 underestimates the true joint probability of the two vectors, because it assumes that corresponding pairs of elements are independent, when some may not be. The mutual information *v* and *u* is then

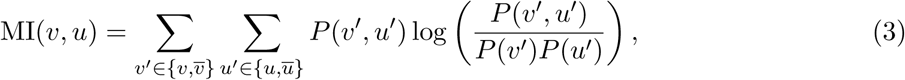

where *v̅* = [1 – *p*_1_,…, 1 – *p*_n_]. Note that, because Equation 2 underestimates the true joint probability of vector v with u, it follows Equation 3 also underestimates the true mutual information of *v* and *u*.

Given a pair of enzymes, the mutual information of their phylogenetic profiles is simply this, according to Equation 3. The mutual information of their evolutionary scenarios is the sum of the mutual information of their respective gain and loss evolutionary scenarios.

### Comparing mutual information to Discrete

Discrete [16] is a method that computes how likely a pair of discrete characters is coevolving in a phylogenetic tree. It takes as input, a phylogenetic tree on a set of objects, and for each of a pair of characters, a vector of discrete states of this character in the objects at the leaves of the tree. It provides a continous-time Markovian substitution model of evolution where the pair of characters evolve independently, and another more complex model where the characters are depedent. Given the input, Discrete allows to compute the log-likelihood of a maximum likelihood (ML) evolutionary scenario under these two models. The ratio of these two log-likelihoods is then an indication of how correlated the evolution of the pair of characters is in the phylogeny. We compute with Discrete this log-likelihood ratio for each pair over the 2732 enzymes listed in S2 Table, given the ultrametric phylogenetic tree depicted in S1 Fig, and compare this score with that of the mutual information of the evolutionary scenarios of the pair.

Discrete takes discrete states on input, and so for each enzyme, we discretize its phylogenetic profile with probability threshold 0.5, *i.e.,* any probability less than 0.5 is set to 0, otherwise it is set to 1. When then compute with Discrete this log-likelihood ratio for each pair of enzymes from randomly selected million of the possible 3 730 546 pairs over the 2732 enzymes, based on their discretized profiles and the ultrametric phylogenetic tree. We then infer (gain and loss) evolutionary scenarios for each enzyme as we did in Evolutionary scenarios, but based instead on the above discretized phylogenetic profile. We then compute, for each pair of enzymes from the above million pairs, the mutual information of the corresponding pair of resulting evolutionary scenarios.

## Community analysis

### The graph

We take the weighted graph *G* = (*V*, *E*) where *V* is the set of our 2732 enzymes, and each edge (*e*, *f*)∈ *E* is weighted by *w*(*e*, *f*) = MI(*e*,*f*). This graph *G* is a complete graph (smallest MI is 3.30e-11). In addition to analyzing *G* itself, we also *threshold* the edges of *G* according to 101 evenly-spaced thresholds *t* ∈ *t_0_*,…,*t*_100_, ranging from smallest MI (*t*_0_ = 3.30e-11) to largest MI (*t*_100_ = 0.104), *i.e.,* for any edge (*e*, *f*) ∈ *E* such that *w*(*e*, *f*) < *t*, we set *w*(*e*, *f*): = 0. Note that if a node *v* of *G* becomes isolated (all edges including *v* have weight zero) in this thresholding process, it is not considered as a member (of the set of nodes) of the resulting graph. When the edges of *G* are thresholded with respect to the threshold *t*, we refer to the resulting graph as *G_t_*, or as “graph *G* at threshold *t*”.

### Community detection

To each of the (101) graphs *G_t_* for the thresholds *t* ∈ *t*_0_,…, *t*_100_, we apply a node-clustering algorithm [3] that finds communities of the nodes of a graph. We also applied a link-clustering algorithm [1] that clusters on the *edges* of the graph, allowing communities (in terms of nodes) to overlap, because a node can appear in more than one edge cluster. All the results for the link commmunities appears in Appendix.

## Communities compared to Kegg modules

The Kegg modules [11] are a collection of manually curated sets of enzymes that participate together in important pathways. We want to compare our communities to these modules. In order to do this, we first downloaded the most recent set of Kegg modules from http://www.genome.jp/kegg/module.html.

Since we are only interested in these modules in terms of the 2732 enzymes of our study, we first restrict these modules to this set of enzymes, remove all duplicate entries from the modules (some modules contain twice the same enzyme), and then discard all modules of size 1. We do this last step because a cluster *(e.g.,* module or community) of size 1 has no meaning when comparing a pair of clusterings of elements: an element is part of a clustering only if it appears paired with some other element in the clustering. Note that this built-in to the community detection algorithms [1,3] – indeed these return communities of size at least 2.

We use a variety of clustering comparison measures to compare our communities to the Kegg modules. In all such measures, a *clustering* of elements from a set *U* of elements is any subset 𝒞 ⊆ 2^*U*^ such that for all *c* ∈ 𝒞 it holds that |*c*| ≥ 2 (as we mentioned above). It hence follows that Kegg modules and our node and link communities are all clusterings on *U*, where *U* is the set of 2732 enzymes of our study – in fact the Kegg modules is also a clustering on the subset *U*′ of 732 enzymes listed in Table. Note that the node communities, for example, have the additional property that for all pairs *c*, *c*′ ∈ 𝒞 it holds that *c* ∪ *c*′ = Ø. The two indexes we consider involve the computation of the following intermediate values for a cluster *c* ∈ 𝒞, with respect to a clustering 𝒞′:

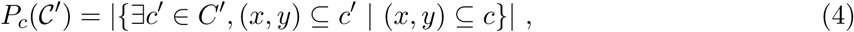

that is, *P_c_*(𝒞′) is the number of pairs from cluster *c* that appear in *some* cluster of clustering 𝒞′. The first index we consider is the *Wallace index*:

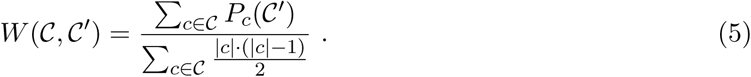

Since the denominator of Equation 5 grows quadratically with the size of custer c, large clusters will dominate this denominator, resulting in smaller Wallace indexes. To more fairly weight the clusters, we propose, as a second index, the *weighted Wallace index*:

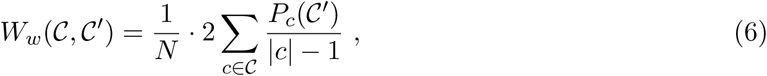

where *N* =Σ_*c*∈𝒞_ |*c*|. We use these three indexes in the following.

Now, since the Kegg modules described above are on 732 enzymes of a possible 2732, in order to compare our communities to the these modules in a meaningful way, we first restrict our communities to the Kegg modules before applying the above indexes. Let *U* be the set of 2732 enzymes of our study (resp., *U*′ be the subset of *U* as listed in Table). Let *C* denote a collection of communities, a clustering on *U* (resp., *C*′ denotes the Kegg modules, a clustering on *U*′). We restrict *C* to the clustering

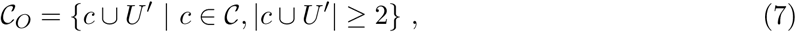

which we refer to as the *U′-overlapping communities,* or simply *overlapping communties,* when the context is clear. We now give some descriptive statistics on the comparision of overlapping communities to Kegg modules.

## Examples of metabolic modules

In addition to comparing to Kegg modules in a rather systematic fashion, we wish to compare our communities to the metabolic modules found in related literature. One of the most related studies to ours is the two works of Yamada *et al.* [20,21], where they find groups of evolutionarily related functions (enzymes) that are also found in similar pathways, which they refer to as *phylogenetic network modules.* We first focus on the modules found in [20].

### Yamada’s Prokaryote Paper [20]

In [20], the authors report a variety of modules found by their method. The largest module they found contains 25 enzymes, and spans both nucleotide and amino acid metabolism, as depicted in Fig. 4 of [20]. The enzymes of this module that span nucleotide and amino acid metabolism are listed in Tables 1 and 2, respectively. Notice that they overlap in the 4 enzymes: 2.1.3.3, 2.1.3.2, 3.5.2.3 and 1.3.98.1.

**Table 1.**
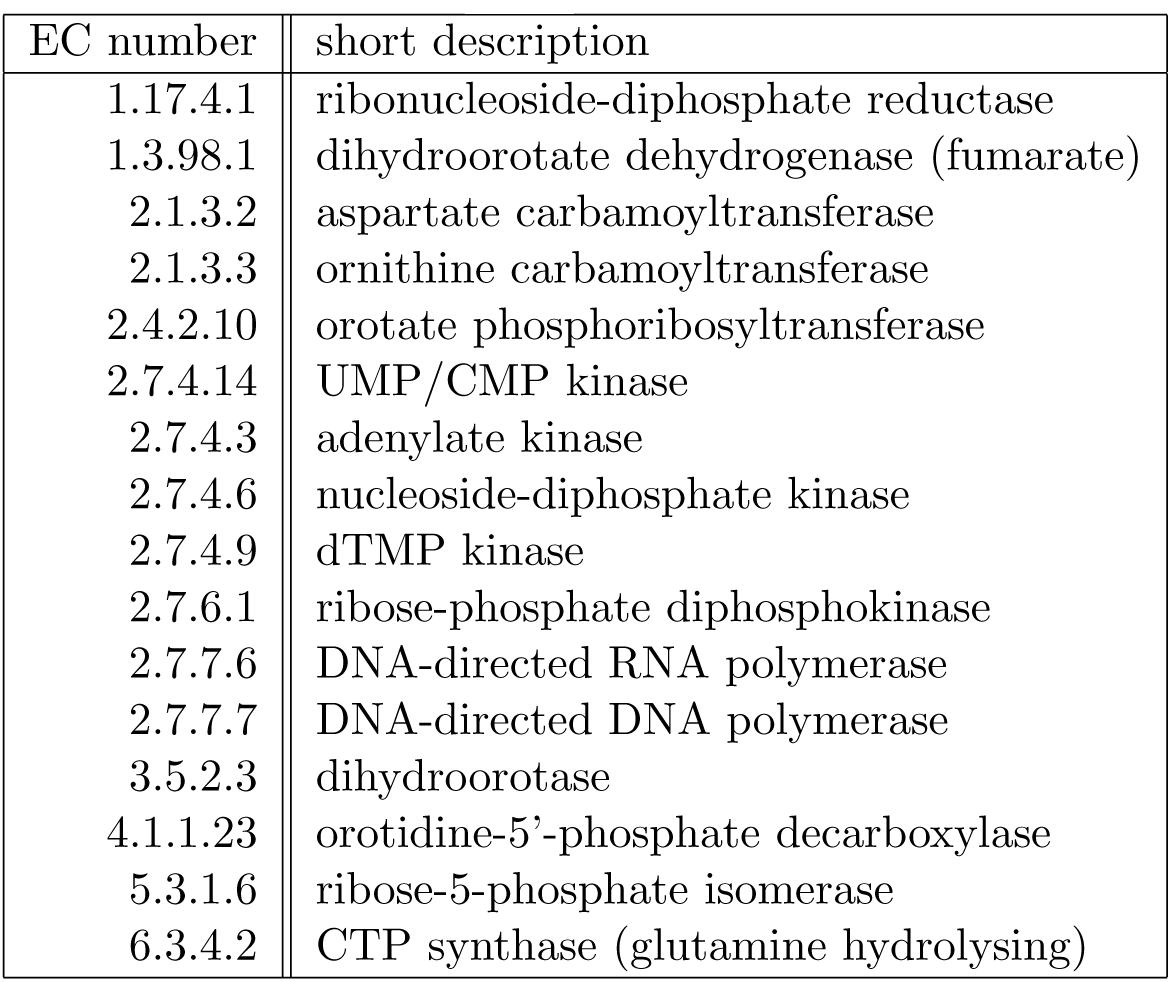
Enzymes of the largest module of [20] that spans nucleotide metabolism, as depicted in Fig. 4 of [20].

Note that enzyme 1.3.3.1 has been renamed to 1.3.98.1 since [20] was published.

Another module found in [20] is one of 7 enzymes that is found in a pathway for peptidoglycan biosynthesis, as depicted in Fig. 5(a) of [20]. These 7 enzymes are listed in Table 3.

The module spanning valine, leucine and isoleucine biosynthesis, depicted in Fig. 5(c) of [20] has the most similarity to our findings, among the modules they found. This modules contains the 6 enzymes listed in Table 4.

A pair of the three modules spanning histidine metabolism, depicted in Fig. 5(d) of [20] has close similarity to our findings as well. The enzymes of these two modules are listed in Tables 5 and 6.

### Yamada’s Eukaryote Paper [21]

In a paper following [20], the same authors apply their method to a wider set of orgaisms, including eukaryotes [21]. Here, they also report a variety of modules found by their method. The largest module they found contains 10 enzymes, and spans the lysine biosythesis pathway and histidine metabolism, as depicted in Fig. 3 of [21]. The enzymes of this module that span the lysine biosynthesis pathway and histidine metabolism are listed in Tables 7 and 8, respectively.

**Table 2.**
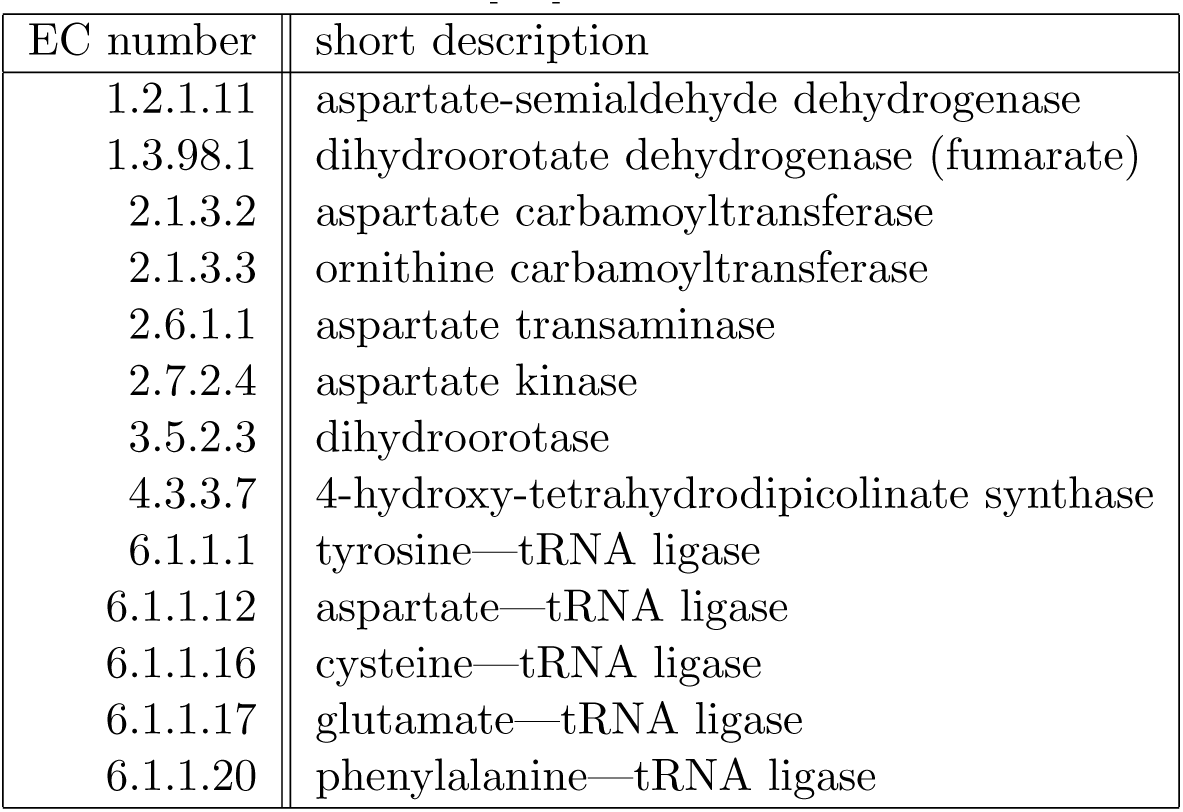
Enzymes of the largest module of [20] that spans amino acid metabolism, as depcited in Fig. 4 of [20].

Note that enzymes 1.3.3.1 and 4.2.1.52 have been renamed to 1.3.98.1 and 4.3.3.7, respectively, since [20] was published.

**Table 3.** Enzymes of a module of [20] that spans a peptidoglycan biosynthesis pathway, as depicted in Fig. 5(a) of [20].

| EC number | short description |
| --- | --- |
| 2.4.1.227 | undecaprenyldiphospho-muramoylpentapeptide beta-N-acetylglucosaminyltransferase |
| 2.7.8.13 | phospho-N-acetylmuramoyl-pentapeptide-transferase |
| 6.3.2.10 | UDP-N-acetylmuramoyl-tripeptide—D-alanyl-D-alanine ligase |
| 6.3.2.13 | UDP-N-acetylmuramoyl-L-alanyl-D-glutamate—2,6-diaminopimelate ligase |
| 6.3.2.4 | D-alanine—D-alanine ligase |
| 6.3.2.8 | UDP-N-acetylmuramate—L-alanine ligase |
| 6.3.2.9 | UDP-N-acetylmuramoyl-L-alanine—D-glutamate ligase |

### Chistoserdova’s Methylotrophy Paper [4]

In [4], the authors study a particular organism and the metabolic functions therein. These functions can be grouped into various modules, which they also report. One module of their study that has some similarity to our findings is the module for the biosynthesis of H_4_F, or tetrahydrofolate, whose enzymes are listed in Table 9.

**Table 4.** Enzymes of a module of [20] that spans a valine, leucine and isoleucine biosynthesis pathway, as depicted in Fig. 5(c) of [20].

| EC number | short description |
| --- | --- |
| 1.1.1.85 | 3-isopropylmalate dehydrogenase |
| 1.1.1.86 | ketol-acid reductoisomerase |
| 2.2.1.6 | acetolactate synthase |
| 2.3.3.13 | 2-isopropylmalate synthase |
| 4.2.1.33 | 3-isopropylmalate dehydratase |
| 4.2.1.9 | dihydroxy-acid dehydratase |

**Table 5.**
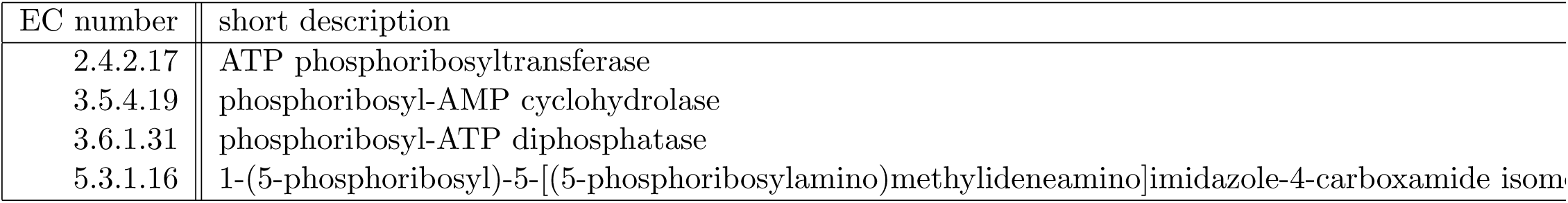
Enzymes of the first module of [20] that spans histidine metabolism, that we compare to, as depicted in Fig. 5(d) of [20].

**Table 6.** Enzymes of the second module of [20] that spans histidine metabolism, that we compare to, as depicted in Fig. 5(d) of [20].

| EC number | short description |
| --- | --- |
| 3.5.2.7 | imidazolonepropionase |
| 3.5.3.8 | formimidoylglutamase |
| 4.2.1.49 | urocanate hydratase |
| 4.3.1.3 | histidine ammonia-lyase |

**Table 7.** Enzymes of the largest module of [21] that spans the lysine biosynthesis pathway, as depicted in Fig. 3 of [21].

| EC number | short description |
| --- | --- |
| 1.1.1.3 | homoserine dehydrogenase |
| 1.2.1.11 | aspartate-semialdehyde dehydrogenase |
| 2.7.2.4 | aspartate kinase |
| 4.3.3.7 | 4-hydroxy-tetrahydrodipicolinate synthase |

Note that enzyme 4.2.1.52 has been renamed to 4.3.3.7 since [21] was published.

Note that enzyme 4.2.1.52 has been renamed to 4.3.3.7 since [21] was published.

**Table 8.** Enzymes of the largest module of [21] that spans histidine metabolism, as depicted in Fig. 3 of [21].

| EC number | short description |
| --- | --- |
| 2.4.2.17 | ATP phosphoribosyltransferase |
| 2.6.1.9 | histidinol-phosphate transaminase |
| 3.5.4.19 | phosphoribosyl-AMP cyclohydrolase |
| 3.6.1.31 | phosphoribosyl-ATP diphosphatase |
| 4.2.1.19 | imidazoleglycerol-phosphate dehydratase |
| 5.3.1.16 | 1-(5-phosphoribosyl)-5-[(5-phosphoribosylamino)methylideneamino]imidazole-4-carboxamide isom |

Note that this module contains the module referred to in Table 5.

Note that this module contains the module referred to in Table 5.

**Table 9.** Enzymes of the module of of [4] that is responsible for the biosynthesis of H4F, as depicted by the hexagonal module of Fig. 1 of [4].

| EC number | short description |
| --- | --- |
| 1.5.1.3 | dihydrofolate reductase |
| 2.5.1.15 | dihydropteroate synthase |
| 2.7.6.3 | 2-amino-4-hydroxy-6-hydroxymethyldihydropteridine diphosphokinase |
| 3.5.4.16 | GTP cyclohydrolase I |
| 3.5.4.29 | GTP cyclohydrolase IIa |
| 4.1.2.25 | dihydroneopterin aldolase |
| 6.3.2.12 | dihydrofolate synthase |
| 6.3.2.17 | tetrahydrofolate synthase |

Note that this module contains the module referred to in Table 5.

## Results

### Genome enzyme assignment with Priam

Fig. 1 illustrates how the probabilities assigned by Priam search of the 2732 specific enzymes of the 1452 organisms of our study are distributed over the unit interval. Fig. 2 illustrates how these 1452 organisms are distributed with respect the number of specific enzymes assigned by Priam to each organism.

If probabilities are *discretized* with theshold 0.5, *i.e.,* enzyme *e* is *present* in an organism if *p*(*e*) ≥ 0.5, otherwise it is absent, then the average number of enzymes present in an organism is 296.78, while the median is 278. The three organisms having the smallest number of enzymes are two bacterial symbionts and a nanoarchaeum: Candidatus Hodgkinia cicadicola Dsem, Nanoarchaeum equitans Kin4-M and Candidatus Carsonella ruddii PV, having respectively 8, 12 and 14 enzymes. The two organisms having the largest number of enzymes present are Homo sapiens (human) and Mus musculus (mouse), having respectively 737 and 725 enzymes.

### Metabolic networks

Fig. 3 illustrates how the number of reactions in the metabolic network (stoichiometry matrix) contstructed by Priam is distributed among the 1452 organisms of our study. The average number of reactions in a metabolic network is 476.3, while the median is 448.5. The three organisms having the smallest number of reactions is the same as the three having the smallest number of enzymes present, but ordered differently: Nanoarchaeum equitans Kin4-M, Candidatus Hodgkinia cicadicola Dsem and Candidatus Carsonella ruddii PV, having respectively 13, 15 and 23 reactions. The two organisms having the largest number of reactions is also the same as the two having the largest number of enzymes present, but also ordered differently: Mus Musculus and Homo sapiens, having respectively 1276 and 1261 reactions.

### Evolutionary scenarios

?? illustrates the gain while ?? illustrates the loss activity on the branches of the species phylogeny with respect to the expected number of gains and losses^1^ of each enzyme.

Fig. 4 depicts the distribution of the probabilites of the ECs at the root of the phylogeny. [**to do**: *something more about bbpancestor*].

### Mutual information

Fig. 5 illustrates how mutual information is distributed among these pairs of enzymes. The highest MI of 8.57e-2 is between Urocanate hydratase (4.2.1.49) and Histidine ammonia-lyase (4.3.1.3), while the lowest MI of 6.75e-4 is between D-malate dehydrogenase (1.1.1.83) N-acylneuraminate cytidylyltransferase (2.7.7.43). [**note:** *more descriptive stats here as well?*]

### Discrete

Fig. 6 is a scatter plot of these *L_D_* – *L_I_* from Discrete against MI for the 952 627 pairs 𝒫. [**to do**: *table of run-time comparisons*] [**note:** *more descriptive stats here? an R^2^ of this plot perhaps?*]

### Phylogenetic profiles

Fig. 7 is a scatter plot of the (species-phylogeny-aware) mutual information of the evolutionary scenarios, against the mutual information of purely the phylogenetic rofiles (as described above).

### Community analysis

#### The graph

Inspecting Fig. 5, one sees that it follows an exponential distribution for the very low values of MI, and then it departs from this (around MI = 0.014). We refer to this threshold, which corresponds to *t*_14_ of the range *t*_0_,…, *t*_100_ from above, as the threshold when MI is *significant,* or simply the *significance threshold.* See Appendix for more details on the selection of this threshold. Fig. 9 illustrates the number of nodes and edges in graph *G_t_*, as a function of threshold *t*. Table 10 is a table of values from the previous figure, for selected values of *t*.

**Table 10.** The number of nodes and edges in *G_t_* for selected values of *t*.

| threshold | vertices | edges | threshold | vertices | edges |
| --- | --- | --- | --- | --- | --- |
| 5 | 1122 | 21848 | 55 | 69 | 70 |
| 10 | 673 | 3413 | 60 | 59 | 57 |
| 15 | 407 | 1359 | 65 | 50 | 44 |
| 20 | 299 | 822 | 70 | 45 | 38 |
| 25 | 235 | 518 | 75 | 32 | 27 |
| 30 | 194 | 385 | 80 | 25 | 22 |
| 35 | 158 | 283 | 85 | 20 | 18 |
| 40 | 131 | 218 | 90 | 12 | 8 |
| 45 | 103 | 141 | 95 | 7 | 4 |
| 50 | 83 | 87 | 100 | 2 | 1 |

The columns are threshold, number of vertices and number of edges in the graph, respectively.

#### Community detection

Fig. 10 illustrates the number of enzymes in the communities *(i.e.,* number of nodes in graph *G_t_*), the number of node communities and the maximum node community size as a function of threshold *t*. Table 11 is a table of values from the previous figure, for selected values of *t*.

#### Kegg modules

After these filtering steps, we are left with 211 modules on 732 enzymes (of a possible 2732), the smallest module is of size 2, while the largest is of size 16. These 732 enzymes are listed in Table. Fig. 11 illustrates the size distribution of the Kegg modules with a histogram. Note that, without any filtering, the Kegg modules contain only 965 enzymes from our set of 2732 of this study, and so this is more a limitation of what is in Kegg, and not our filtering steps.

#### The overlapping communities

Fig. 12 illustrates the number of enzymes in the communities, the number of such communities that are overlapping and the weighted Wallace index, from *G_t_* as a function of threshold *t*. Table 12 is a table of values from the previous figure, for selected values of *t*.

**Table 11.** The number of enzymes, the number of communities and maximum community size from *G_t_* for selected values of *t*.

| thresh. | enzy. | comm. | max size | thresh. | enzy. | comm. | max size |
| --- | --- | --- | --- | --- | --- | --- | --- |
| 5 | 1122 | 38 | 336 | 55 | 69 | 26 | 7 |
| 10 | 673 | 75 | 121 | 60 | 59 | 23 | 7 |
| 15 | 407 | 88 | 45 | 65 | 50 | 20 | 6 |
| 20 | 299 | 86 | 39 | 70 | 45 | 19 | 6 |
| 25 | 235 | 76 | 31 | 75 | 32 | 13 | 6 |
| 30 | 194 | 64 | 26 | 80 | 25 | 10 | 6 |
| 35 | 158 | 55 | 22 | 85 | 20 | 8 | 5 |
| 40 | 131 | 46 | 20 | 90 | 12 | 5 | 3 |
| 45 | 103 | 37 | 12 | 95 | 7 | 3 | 3 |
| 50 | 83 | 32 | 7 | 100 | 2 | 1 | 2 |

The columns are threshold, number of enzymes, number of communities and maximum community size, respectively.

### Examples of metabolic modules

Here, for each module listed in Examples of metabolic modules, we present all its subsets (of size at least two) that are found in the node communities for all of the thresholds *t*.

**Yamada’s Prokaryote Paper [20].**

**Yamada’s Eukaryote Paper [21].**

**Chistoserdova’s Methylotrophy Paper [4].**

**Table 12.**
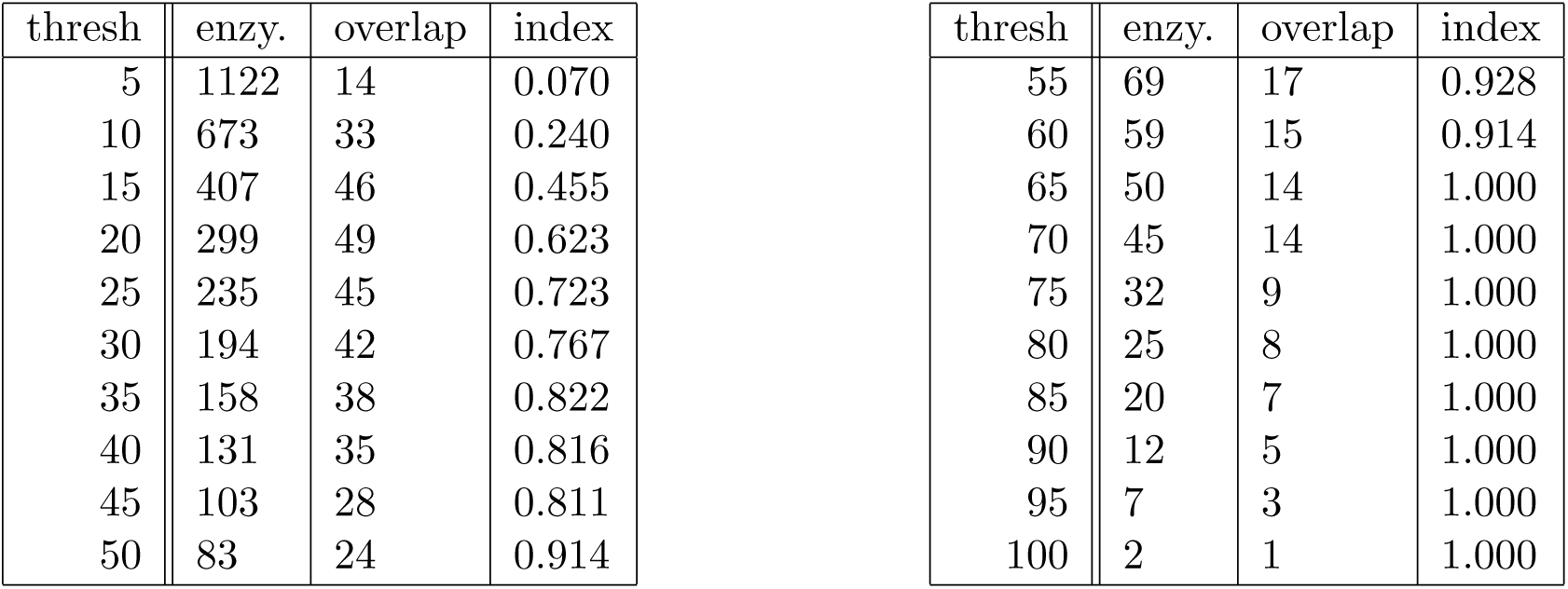
The number of enzymes, the number of overlapping communities and the weighted Wallace index of overlapping communties for selected values of *t*.

The columns are threshold, number of enzymes, number of overlapping communities and the weighted Wallace index of overlapping communities, respectively.

**Table 13.** Containment of the module of Table 1 in the node communities.

|  |  |  |  |  |  |  |  |  |  |  |  |
| --- | --- | --- | --- | --- | --- | --- | --- | --- | --- | --- | --- |
| 2.1.3.2 | 2.1.3.3 | 2.4.2.10 | 2.7.4.3 | 2.7.4.6 | 2.7.4.9 | 2.7.6.1 | 3.5.2.3 | 4.1.1.23 | 6.3.4.2 | + | [1,5] |
| - | 2.1.3.3 | - | - | 2.7.4.6 | - | - | - | - | 6.3.4.2 | +++ | [7] |
| - | 2.1.3.3 | - | - | 2.7.4.6 | - | - | - | - | - | ++++ | [8,10] |
| 2.1.3.2 | - | 2.4.2.10 | 2.7.4.3 | - | - | - | 3.5.2.3 | 4.1.1.23 | 6.3.4.2 | +++ | [8] |
| 2.1.3.2 | - | 2.4.2.10 | - | - | - | - | 3.5.2.3 | 4.1.1.23 | - | +++++ | [10,30] |
| 2.1.3.2 | - | - | - | - | - | - | 3.5.2.3 | - | - | ++++++ | [31,49] |
| - | - | 2.4.2.10 | - | - | - | - | - | 4.1.1.23 | - | ++++++ | [31,39] |
| 2.1.3.2 | - | 2.4.2.10 | - | - | - | - | 3.5.2.3 | 4.1.1.23 | 6.3.4.2 | ++++ | [9] |
| 2.1.3.2 | 2.1.3.3 | 2.4.2.10 | - | 2.7.4.6 | 2.7.4.9 | - | 3.5.2.3 | 4.1.1.23 | 6.3.4.2 | ++ | [6] |
| 2.7.4.14 | 2.7.4.9 | 5.3.1.6 |  |  |  |  |  |  |  | + | [7] |

In the first column, each row contains a (sub-) set of enzymes (by EC code) from the module that are found in a node community for the range(s) of thresholds t of the third column. Sets in rows that are grouped by a single box (not separated by horizontal lines) are nested, as indicated by the second column: any set in a row with more +’s that appears in a row below another set is a subset of that set.

**Table 14.**
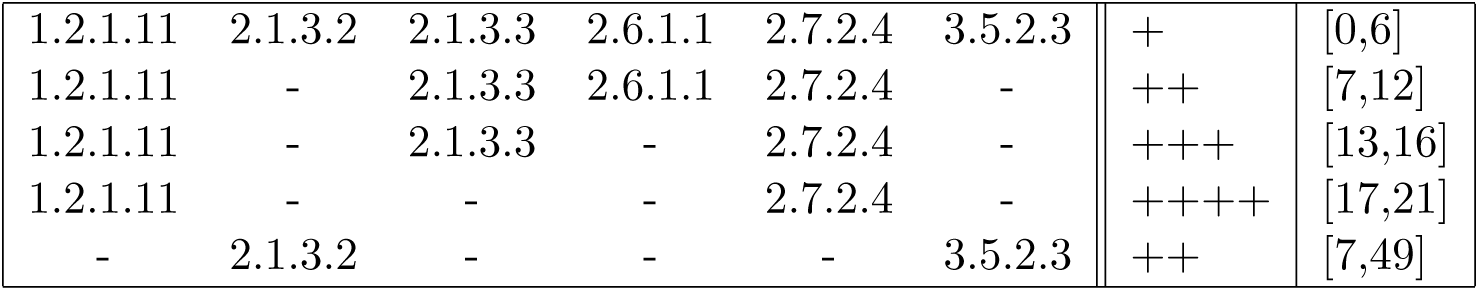
Containment of the module of Table 2 in the node communities.

See legend of Table 13

**Figure 1.**
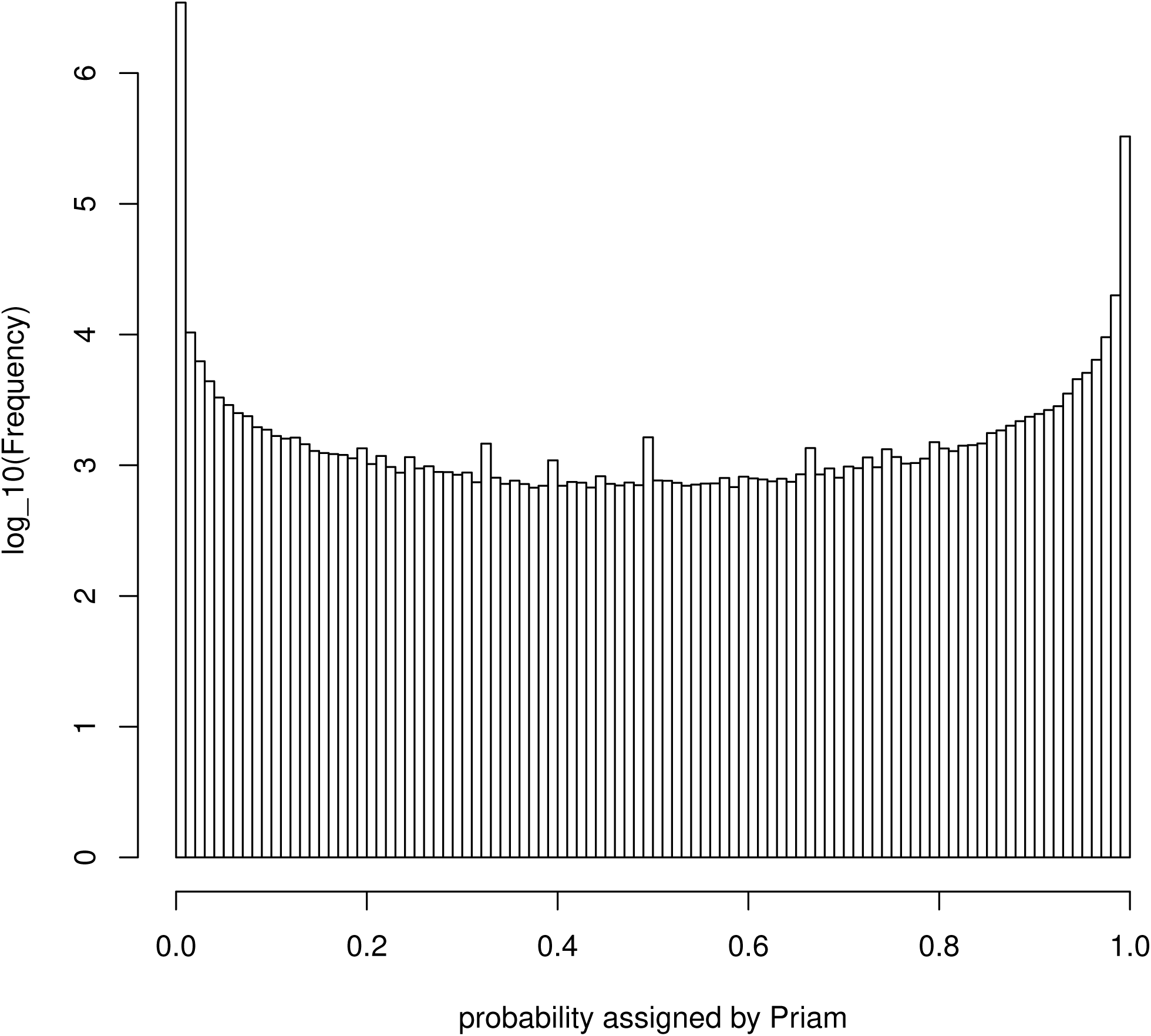
Distribution of probabilities assigned by Priam (search) to the enzymes of the organisms. Histogram of the (3 966 864) probabilities over the 2732 specific enzymes and 1452 organisms of our study.

**Table 15.**
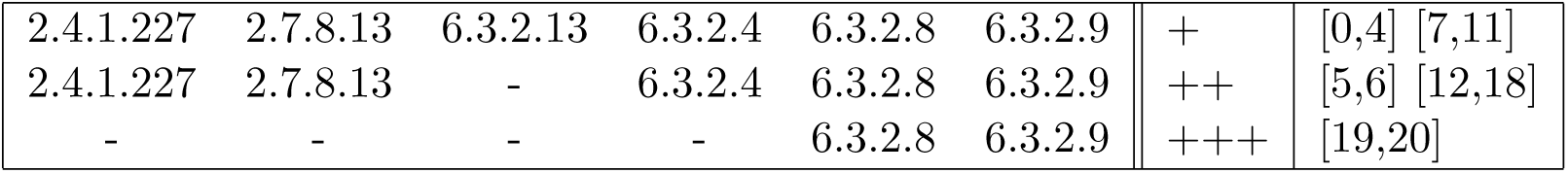
Containment of the module of Table 3 in the node communities.

See legend of Table 13

**Figure 2.**
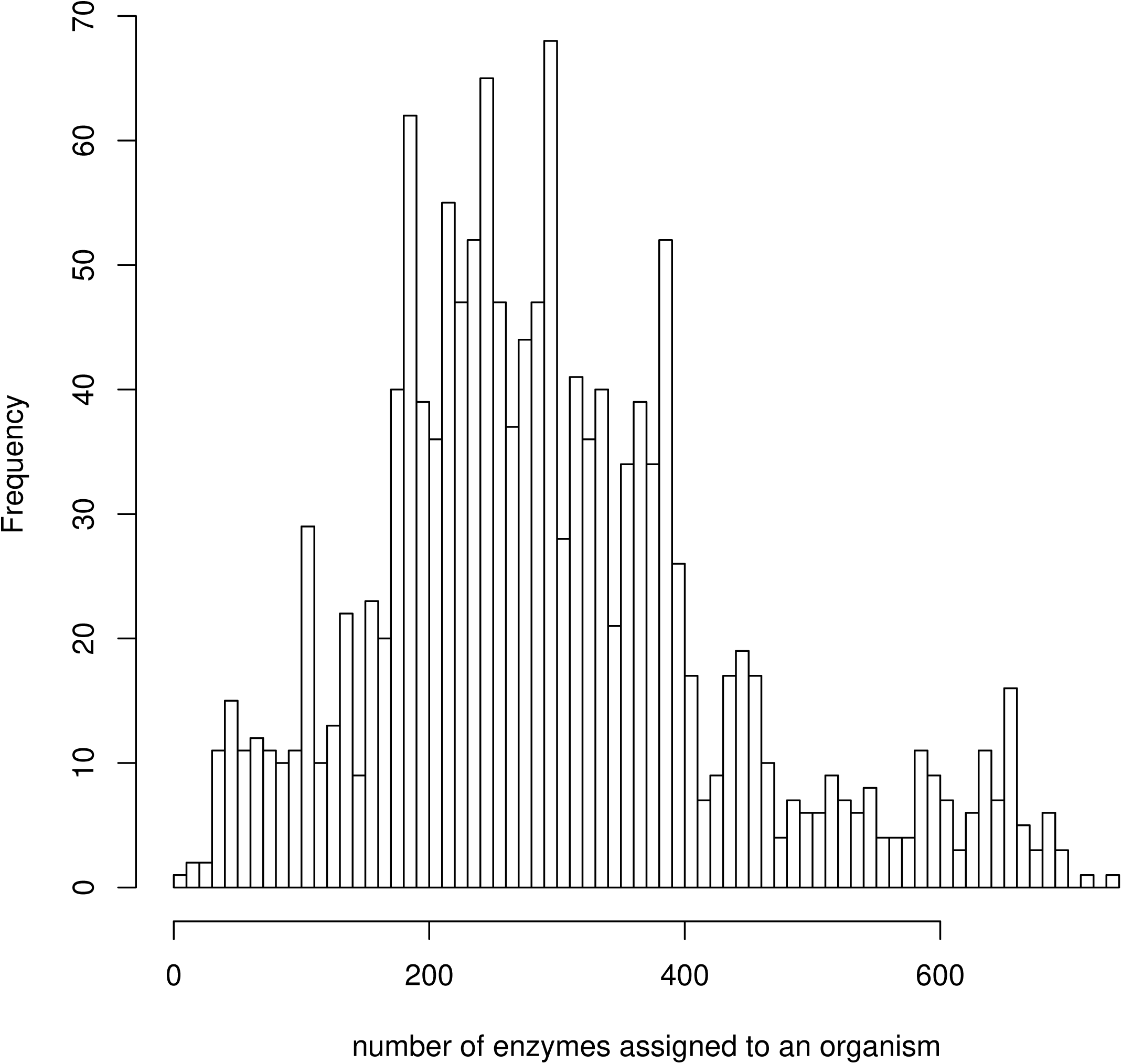
Distribution of organisms with respect to number of enzymes assigned by Priam. Histogram of the sum over the vector of (2732) probabilities of (specific) enzyme presence of each organism given by Priam, for the 1452 organisms of our study.

**Figure 3.**
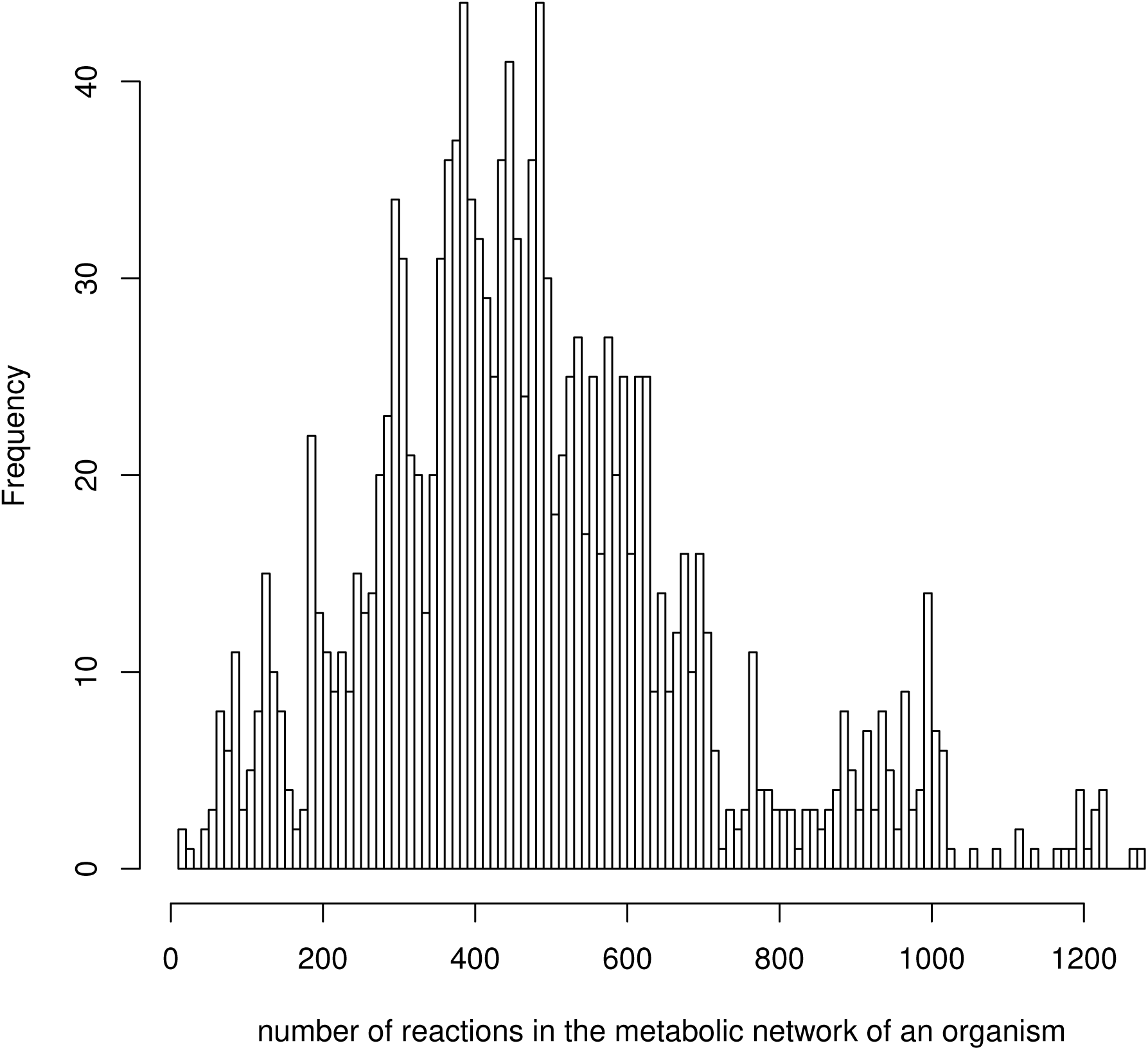
Distribution of organisms with respect to number of reactions in its metabolic network. Histogram of number of reactions in the metabolic network constructed by Priam search of each organism, for the 1452 organisms of our study.

**Table 16.** Containment of the module of Table 4 in the node communities.

|  |  |  |  |  |  |  |  |
| --- | --- | --- | --- | --- | --- | --- | --- |
| 1.1.1.85 | 1.1.1.86 | 2.2.1.6 | 2.3.3.13 | 4.2.1.33 | 4.2.1.9 | + | [0,61] |
| 1.1.1.85 | - | - | 2.3.3.13 | 4.2.1.33 | - | ++ | [62,69] |
| 1.1.1.85 | - | - | 2.3.3.13 | - | - | +++ | [70,72] |
| - | 1.1.1.86 | 2.2.1.6 | - | - | 4.2.1.9 | ++ | [62,67] |
| - | 1.1.1.86 | - | - | - | 4.2.1.9 | +++ | [68,72] |

See legend of Table 13

**Figure 4.**
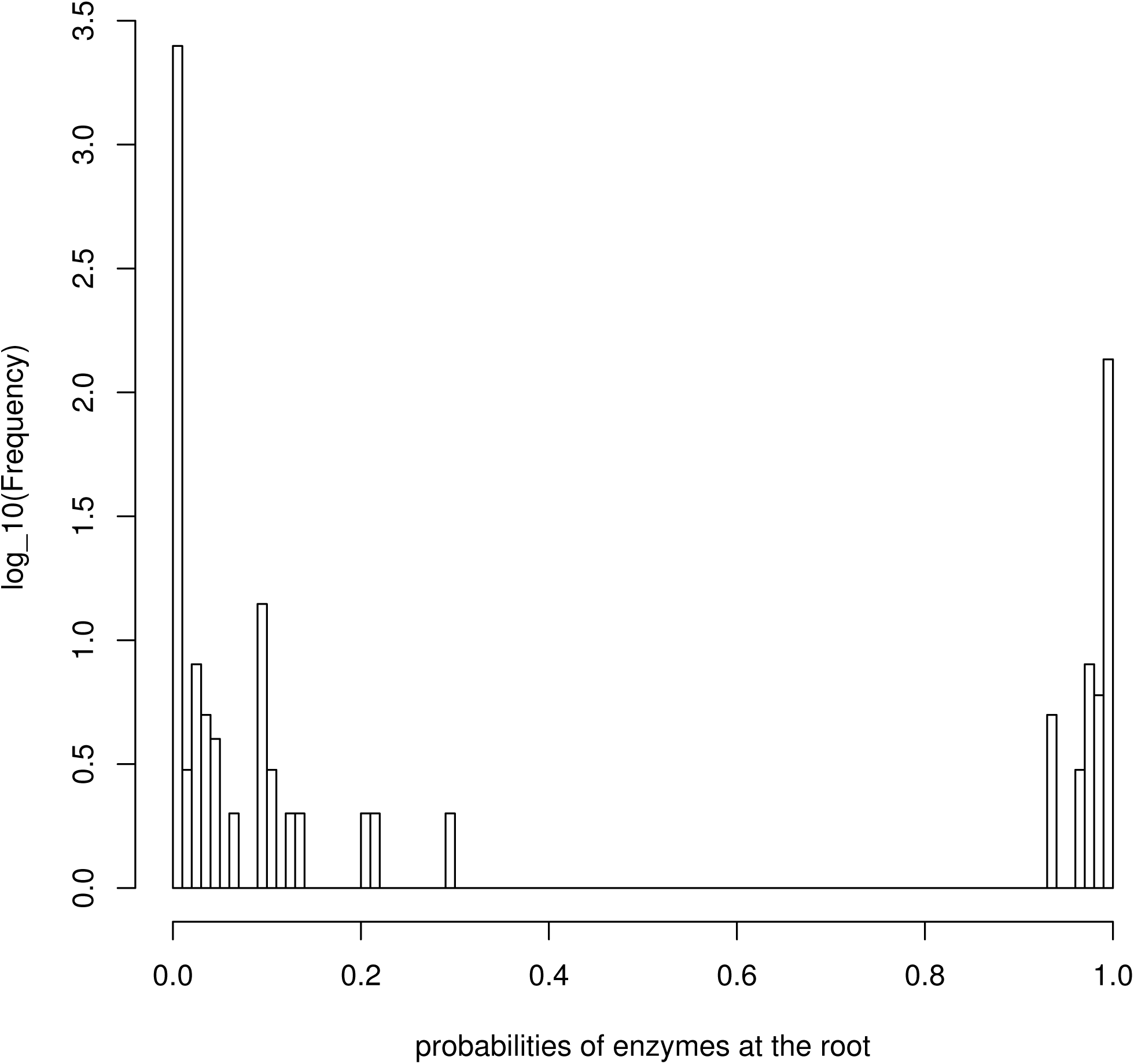
Distribution of probabilities of the enzymes at the root of the species phylogeny. Histogram of the probabilities of the 2732 enzymes at the common ancestor of all species in (the root of) the species phylogeny.

**Table 17.** Containment of the module of Table 5 in the node communities.

|  |  |  |  |  |  |
| --- | --- | --- | --- | --- | --- |
| 2.4.2.17 | 3.5.4.19 | 3.6.1.31 | 5.3.1.16 | + | [0,81] |
| 2.4.2.17 | 3.5.4.19 | - | 5.3.1.16 | ++ | [82,87] |
| 2.4.2.17 | 3.5.4.19 | - | - | +++ | [88,92] |

See legend of Table 13

**Figure 5.**
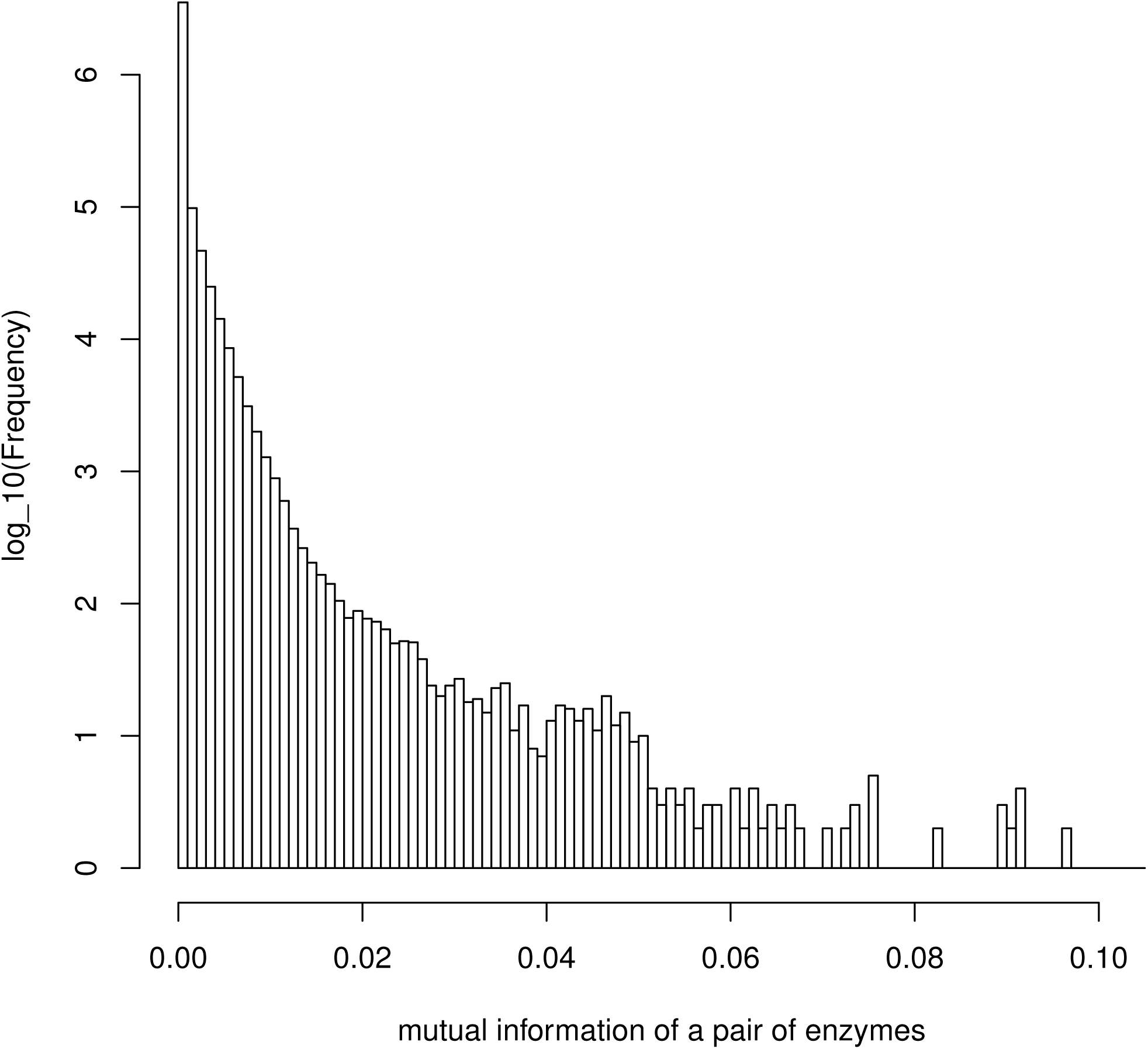
Distribution of mutual information with respect to enzyme pairs. Histogram of mutual information of the 3 730 546 pairs over the 2732 enzymes of our study.

**Table 18.** Containment of the module of Table 6 in the node communities.

|  |  |  |  |  |  |
| --- | --- | --- | --- | --- | --- |
| 3.5.2.7 | 3.5.3.8 | 4.2.1.49 | 4.3.1.3 | + | [0,23] |
| 3.5.2.7 | - | 4.2.1.49 | 4.3.1.3 | ++ | [24,99] |
| - | - | 4.2.1.49 | 4.3.1.3 | +++ | [100] |

See legend of Table 13

**Figure 6.**
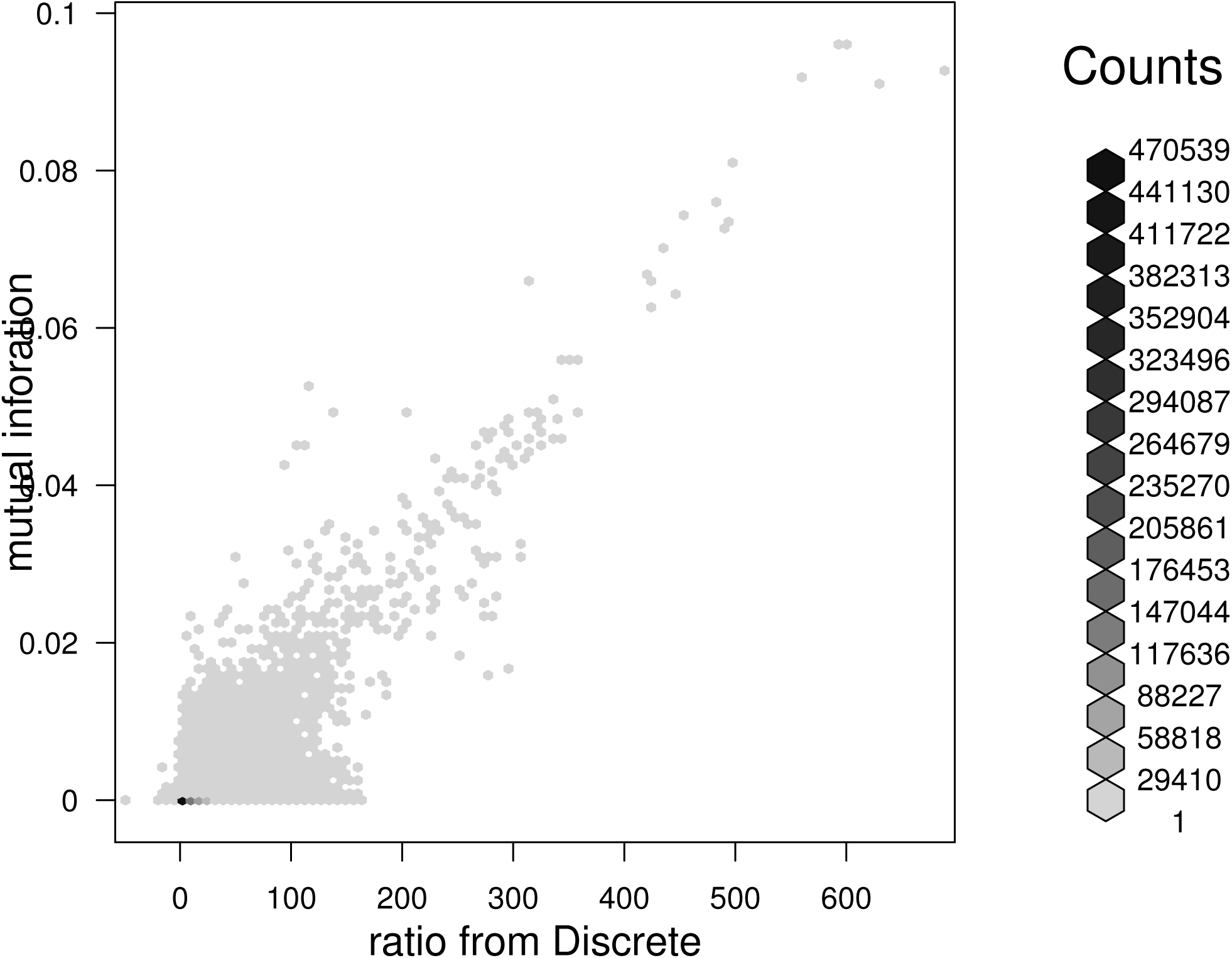
Discrete versus mutual information. Scatter plot of the log-likelihood ratio *L_D_* – *L_I_* from Discrete against mutual information for the 952 627 pairs 𝒫.

**Table 19.** Containment of the module of Table 7 in the node communities.

|  |  |  |  |  |
| --- | --- | --- | --- | --- |
| 1.1.1.3 | 1.2.1.11 | 2.7.2.4 | + | [0,16] |
| - | 1.2.1.11 | 2.7.2.4 | ++ | [17,21] |

See legend of Table 13

**Table 20.** Containment of the module of Table 8 in the node communities.

|  |  |  |  |  |  |  |  |
| --- | --- | --- | --- | --- | --- | --- | --- |
| 2.4.2.17 | 2.6.1.9 | 3.5.4.19 | 3.6.1.31 | 4.2.1.19 | 5.3.1.16 | + | [0,63] |
| 2.4.2.17 | - | 3.5.4.19 | 3.6.1.31 | 4.2.1.19 | 5.3.1.16 | ++ | [64,81] |
| 2.4.2.17 | - | 3.5.4.19 | - | 4.2.1.19 | 5.3.1.16 | +++ | [82,87] |
| 2.4.2.17 | - | 3.5.4.19 | - | 4.2.1.19 | - | ++++ | [88,92] |
| - | - | 3.5.4.19 | - | 4.2.1.19 | - | +++++ | [93,94] |

See legend of Table 13

**Figure 7.**
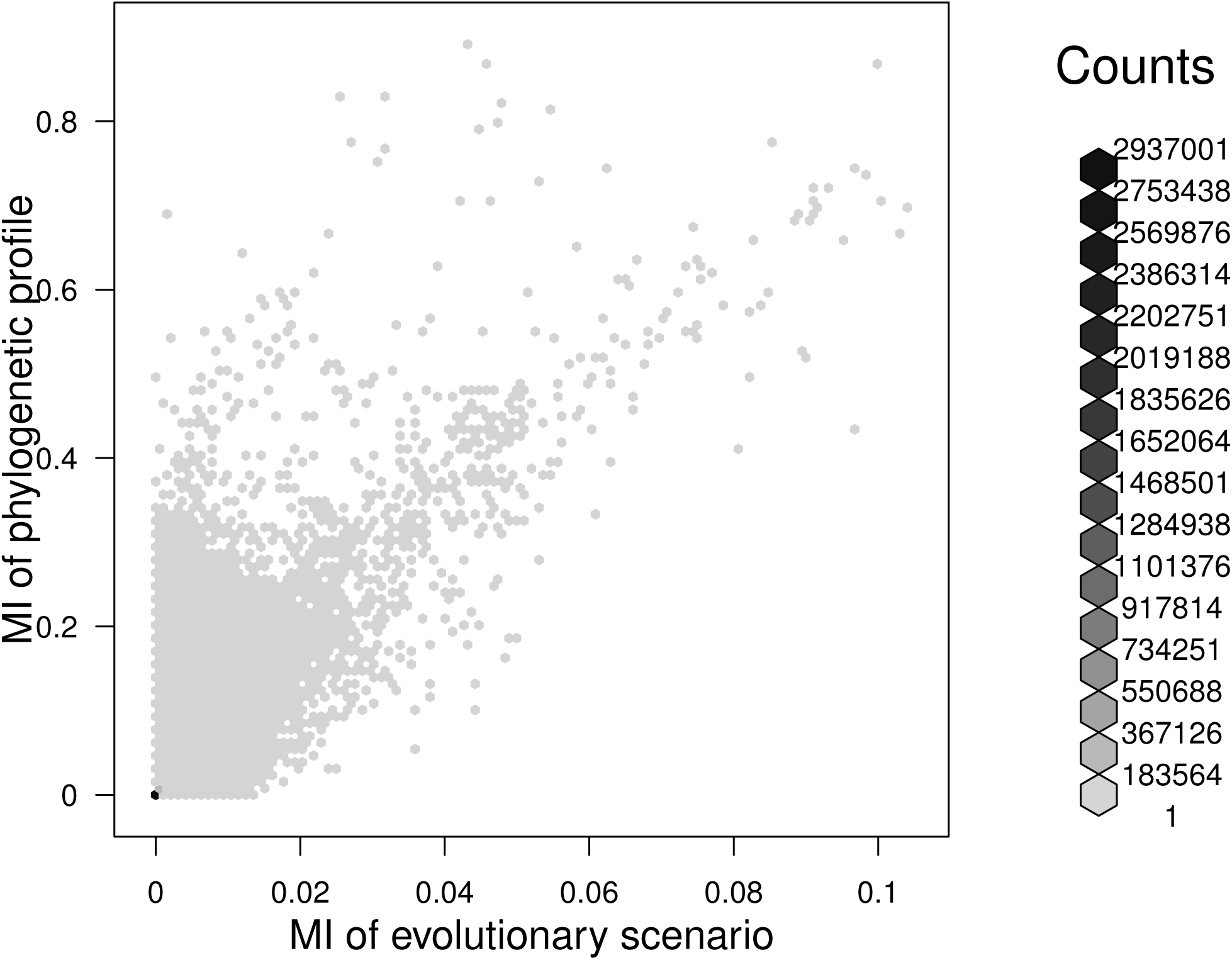
Mutual information of the evolutionary scenarios versus mutual information of the phylogenetic profiles. Scatter plot of the mutual information of the evolutionary scenarios against mutual information of purely the phylogenetic profiles for all pairs of the 2732 enzymes of our study.

**Table 21.** Containment of the module of Table 9 in the node communities.

|  |  |  |  |  |  |  |
| --- | --- | --- | --- | --- | --- | --- |
| 2.5.1.15 | 2.7.6.3 | 3.5.4.16 | 3.5.4.29 | 4.1.2.25 | + | [0,4] |
| 2.5.1.15 | 2.7.6.3 | 3.5.4.16 | - | 4.1.2.25 | ++ | [5,9] [12] |
| 2.5.1.15 | 2.7.6.3 | - | - | 4.1.2.25 | +++ | [10,11] [13,17] |
| 2.5.1.15 | 2.7.6.3 | - | - | - | ++++ | [18,43] |
| 6.3.2.12 | 6.3.2.17 |  |  |  | + | [2,4] |

See legend of Table 13

**Figure 8.**
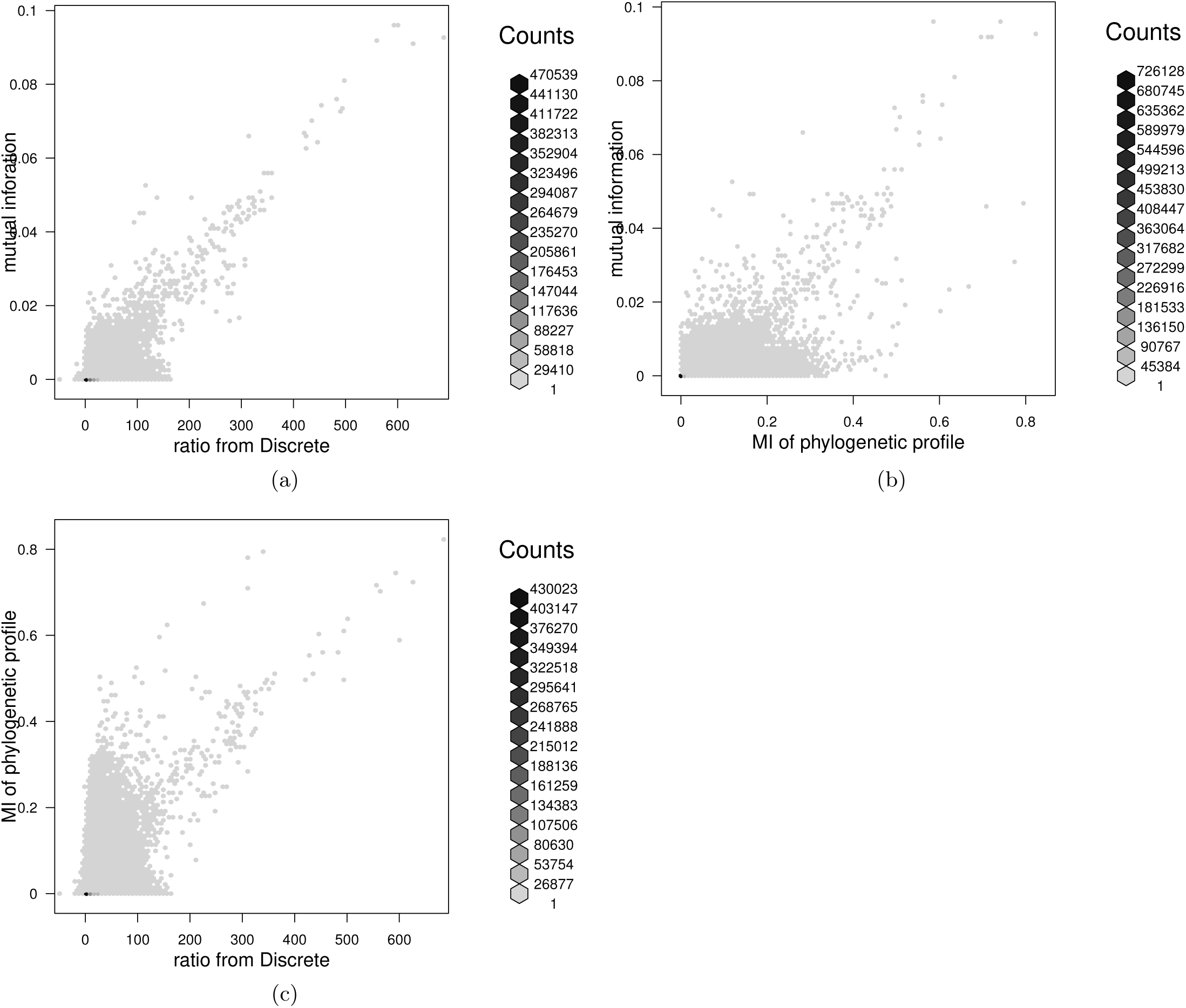
A comparative view of mutual information of evolutionary scenarios, Discrete and mutual information of phylogenetic profiles. For the 952 627 pairs P, we give scatter plots of (a) the log-likelihood ratio *L_D_* – *L_I_* from Discrete against mutual information of evolutionary scenarios, (b) the mutual information of phylogenetic profiles against mutual information of evolutionary scenarios and (c) *L_D_* – *L_I_* of Discrete against mutual information of phylogenetic profiles, in a convenient format that allows the comparison and contrast of the three. Note that, to fairly compare, phylogenetic profiles are discretized using probability threshold 0.5, and the evolutionary scenarios (from which MI is computed) are those obtained by said discretized phylogenetic profiles.

**Figure 9.**
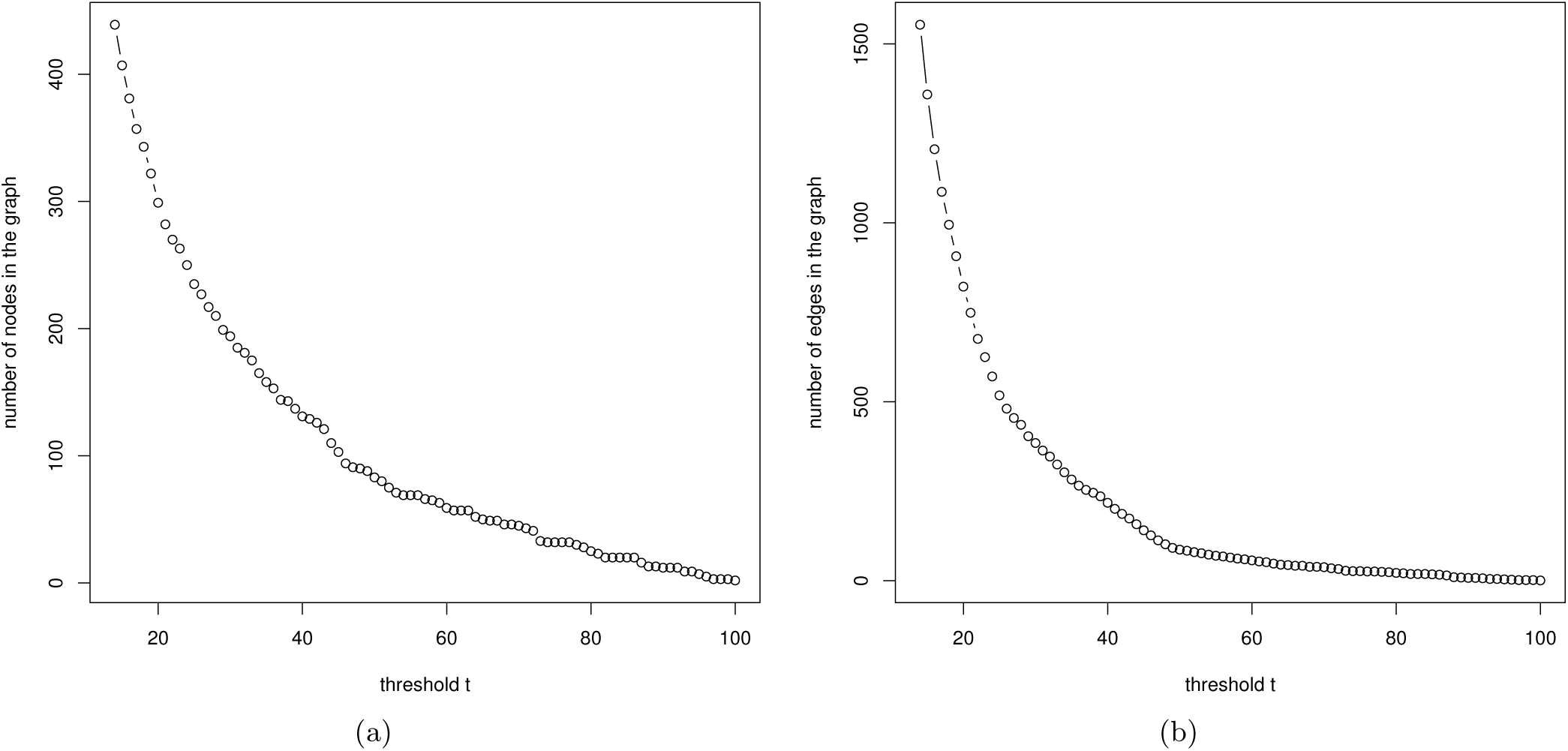
The number of nodes and edges in graph *G_t_* as a function of threshold *t*. The number of (a) nodes in *G_t_* and (b) edges in *G_t_* for the range *t*_14_,…, *t*_100_ of threholds *t*.

**Figure 10.**
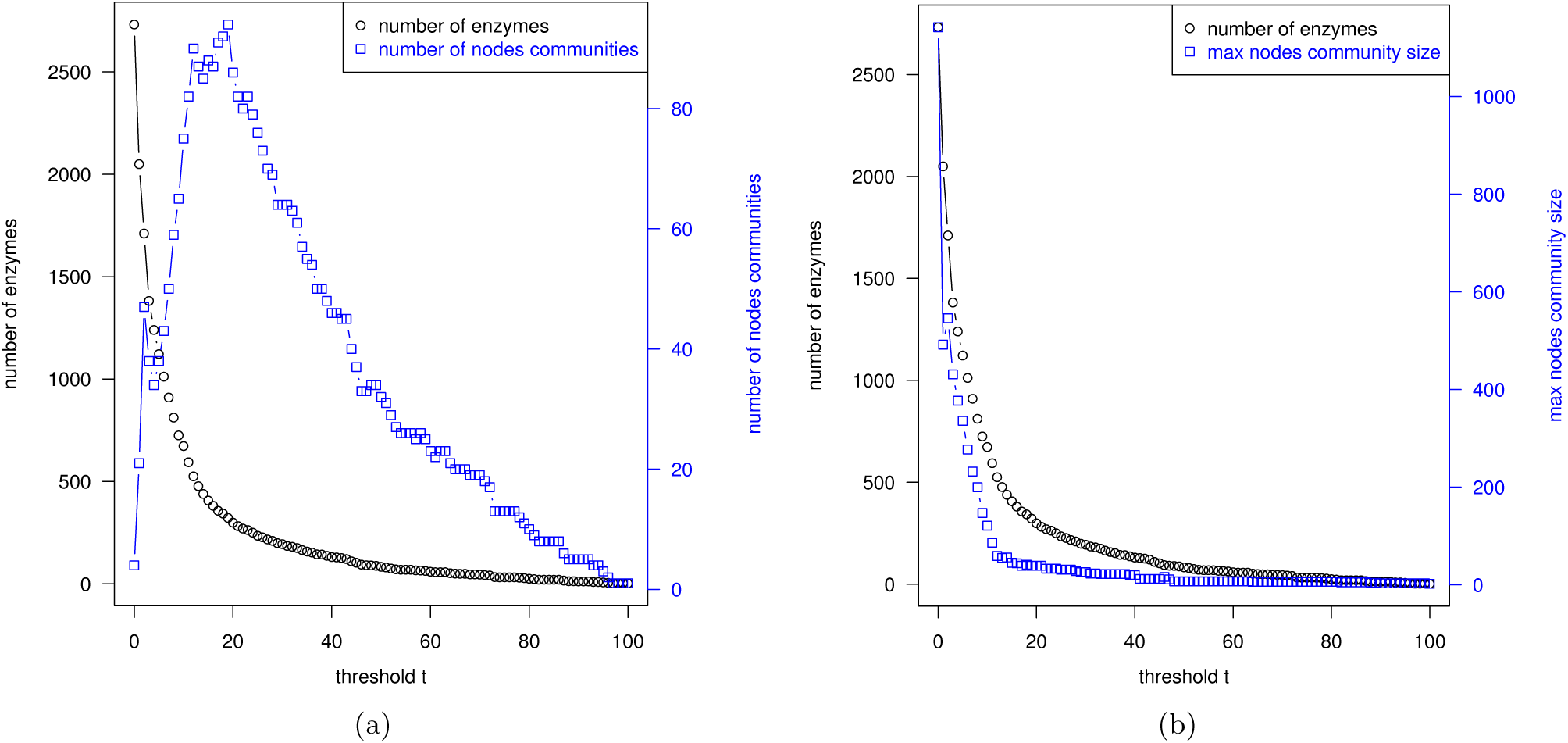
The number of enzymes, the number of communities and the maximum community size from graph *G_t_* as a function of threshold *t*. The number of enzymes, (a) the number of communities and (b) the maximum size of a community from *G_t_* as a function of *t*.

**Figure 11.**
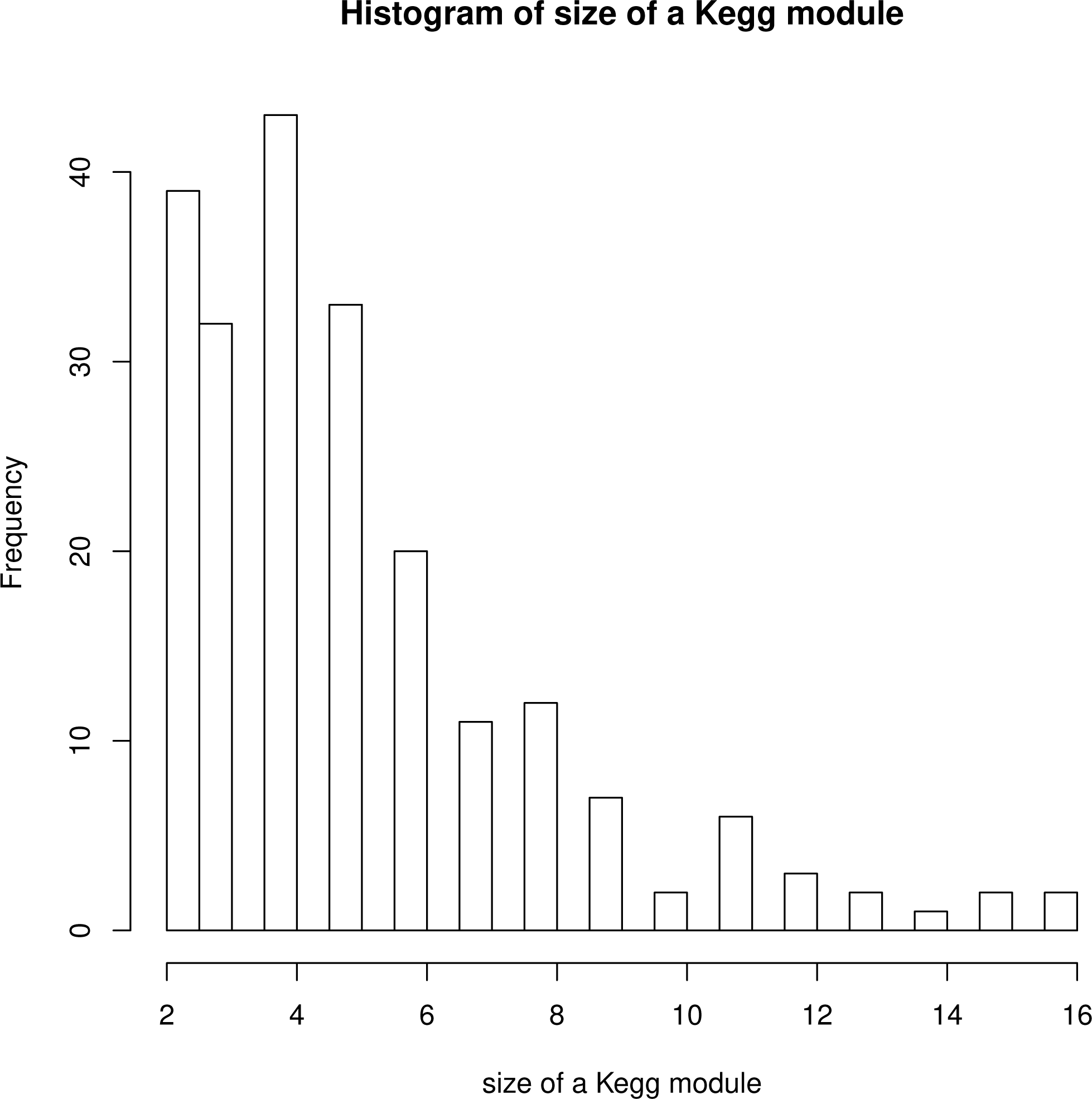
Distribution of the sizes of the Kegg modules. Histogram of the sizes of the Kegg modules restricted to modules of size two containing unique enzymes after restricting to the 2732 enzymes of our study.

**Figure 12.**
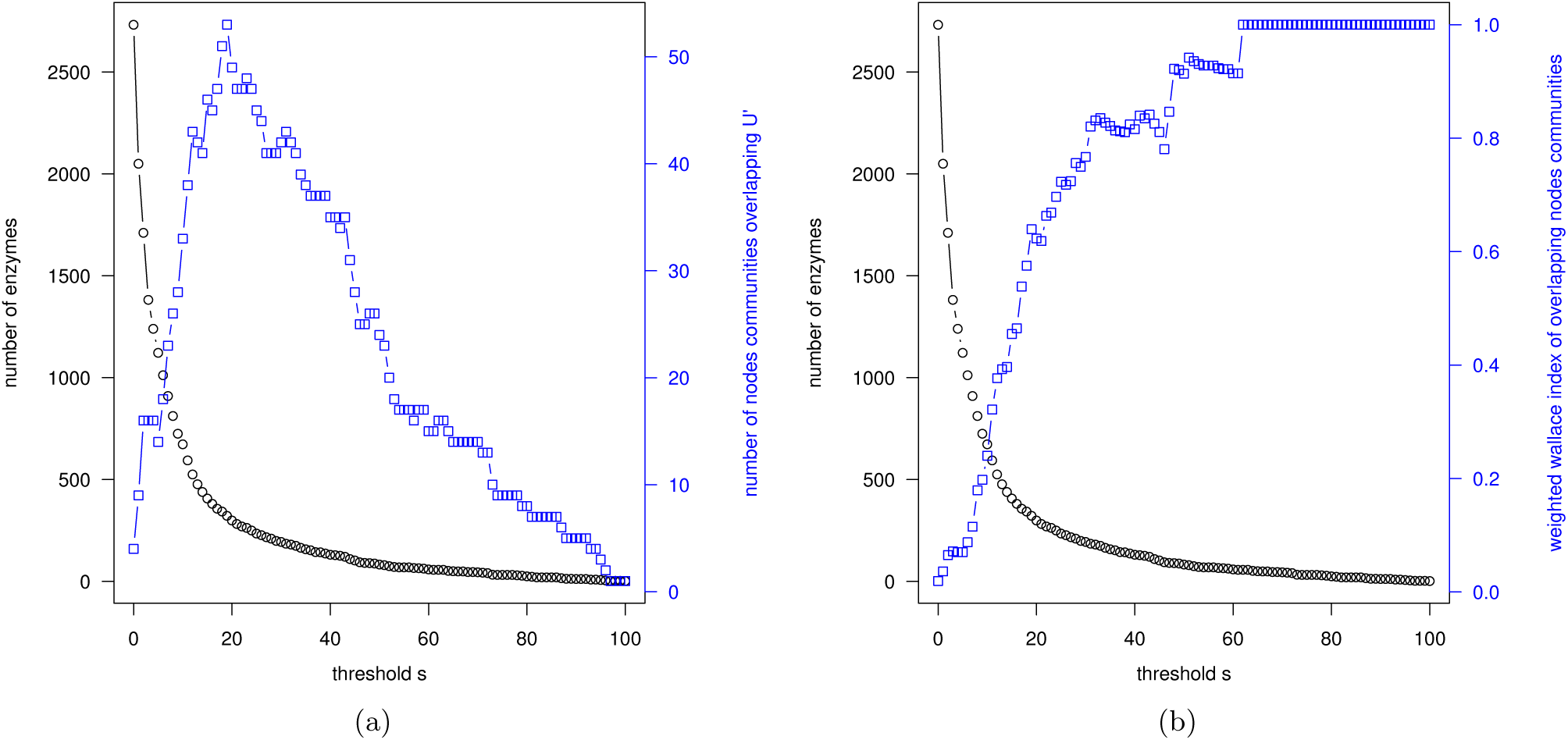
The number of enzymes, (a) the number of overlapping communities and (b) the weighted Wallace index of overlapping communities from *G_t_* as a function of threshold *t*.

## Discussion

### Genome enzyme assignment: a probabilistic framework

Previous related works [12,20,21] also deal with phylogenetic profiles, however these are obtained using orthology data from the KEGG database [10,11]. KEGG orthology data is binary (1: present, or 0: absent) data on the enzymes in an organism, which is less general than the probabilistic framework we present above. Moreover, this type of approach is constrained to the manually curated orthogy data in KEGG. PRIAM, being more automated – delivering enzyme probabilities based purely on protein sequence content given as input – allows the flexibility of studying any set of organisms one desires, as we have done with the HOGENOM [17] organisms.

### Evolutionary scenarios: taking phylogeny into account

Of course, in turn, reasoning about correlatedness with vectors of branch probabilites now treats each branch as being independent, in terms of the gain and loss events that occur [**to do:** *maybe back-up somehow why this independence assumption is not so serious?*]. One thing this is certain is that these vectors of branch probabiities are not biased by phylogenetic inertia [**to do:** *back this up too?*].

### Correlated evolutionary scenarios: mutual information

The most important reason for choosing MI is that the MI shared by a pair of vectors can only be as high as the amount of information, or *entropy* [19], contained in a single vector (see Materials and Methods). Universal enzymes will have a low entropy (see Materials and Methods) and, hence, will not have significantly high MI. Conversely, with measures such as JC or Pearson correlation, universal enzymes pose a problem as these measures tend to 1 (the maximum possible correlation in the two measures) as both enzymes in the pair tend to universality [**to do:** *statter plot depicting this*]. Two enzymes that both happen to be universal are not necessarily correlated, and so the presence of universal enzymes bias the distribution of correlation among pairs to the high end. In order to cope with this, most methods filter out such universal enzymes, as is done in [12]. This, and other advantages of MI over other statisitical correlations is studied in [13].

### Evolutionary scenarios: taking phylogeny into account

One could then use directly these vectors of probabilities of enzyme presence in order to infer the degree to which pairs of enzymes are correlated in a similar way that is done in methods that use *phylogenetic profiles* [12,20,21]. However, this ignores completely the phylogeny, treating each species as being independent, in terms of the presence of the pair of enzymes in question. This is far from being the case, since some species are much more phylogenetically realated than others. This bias has been known for decades, and has been referred to in the literature as *phylogenetic inertia* [2], a term we will use here to summarize this concept. Here, since we do have a species phylogeny at our disposal (see ??), we leverage its information to avoid this bias of phylogenetic inertia. We reason with these vectors of probabilites on the branches of the tree to infer the degree to which pairs of enzymes are correlated, rather than simply probabilities at the leaves.

### Mutual information

Given a pair of enzymes, *i.e.,* the associated pair of vectors of branch probabilities, or phylogenetic profiles, there is a variety of measures one could employ to infer the degree to which this pair is correlated. Examples include statistical correlation functions, such as Pearson correlation, as used in [12], or the *Jaccard Coefficient* (JC) (see [9] and Materials and Methods) as used in [20,21]. In our work, we choose to use the *mutual information* (MI) shared by the pair of vectors (see Materials and Methods for details). The notion of MI comes from the area of information theory [19], and has had much use in areas such as information retrieval [14] and signal processing [8].

### Discrete

The suite of programs *Bayestraits,* developed by Mark Pagel [16], has a subroutine that can be used to infer the degree to which a pair of discrete characters, *e.g.,* enzymes, is co-evolving in a phylogeny (see Materials and Methods for details). Since we only consider this single subroutine, here we refer to it simply as *Bayestraits.* Since, as in our approach, Bayestraits also takes into account the phylogeny, we compared mutual information to what is obtained with Bayestraits. Since Bayestraits takes only *discrete* characters as input, we *discretized* our vectors of enzyme presence probabilities with a probability threhold of 0.5, *i.e.,* for each element *i* of the a vector, if *p*(*i*) < 0.5 we set it to 0 (absent), otherwise we set it to 1 (present). Since Bayestraits is quite time expensive, we then ran Bayestraits on one million randomly selected pairs (of the 3 730 546 pairs) of enzymes. We then computed the mutual information of pairs of enzymes in terms of these discretized vectors (see Materials and Methods for details). Fig. 6 depicts the relationship between Bayestraits and mutual information in the form of a scatter plot.

As a proof of concept – why it is important to take into account the phylogeny for inferring correlatedness of pairs of enzymes – we also compared the mutual information directly of enzyme presence vectors to what is obtained with Bayestraits. Fig ?? illustrates the values by Bayestraits plotted against the mutual information of discretized enzyme presence probability vectors with a probablity threshold of 0.5 for the same million randomly selected pairs as in Fig. 6

One drawback of Discrete is that it can support only discrete states, unlike MapNH, which also supports continuous (probabilistic) states. In order to compare the two on the same grounds, we use as inputs to the two methods: the phylogenetic profiles of the 2732 enzymes of our study, discretized with probability threshold 0.5, along with the species phylogeny. Another drawback of Discrete is that it is more time expensive than our MapNH/MI pipeline, and hence, of the 3 730 546 possible pairs over the 2732 enzymes of our study, we randomly select one million such pairs for analysis. For each of the 2732 enzymes of our study, we input its phylogenetic profile, discretized according to probability threshold 0.5, along with the species phylogeny into MapNH to infer gain and loss evolutionary scenarios of each enyzme. We then computed the MI of each pair over the 2732 enzymes, with respect to these evolutionary scenarios based on organism presence vectors. Then, for Discrete, for each pair of the randmly selected million, we input its corresponding pair of discretized phylogenetic profiles and the species phylogeny into Discrete to obtain this ratio *L_D_* – *L_I_*. Of this million, 952 627 pairs succeeded – failure to succeed occurred due to either inability of the method to find a set of starting parameters, or the method not terminating for a pair after 30 minutes^2^. [**to do:** *say something about the organism presence vectors that are all-zero].* We refer to this set of 952 627 pairs throughout the paper as 𝒫.

### Summary

In summary, our framework offers three advantages over similar studes [12,20,21], in that (a) it deals with probabilities, which is more general; (b) takes into account the phylogeny with branch probabilities, mitigating the bias of phylogenetic inertia; and (c) infers correlation using MI, which does not give too much weight to universal reactions. There is another well-known method for assessing the degree to which pairs of characters (enzymes in our case) are co-evolving, that has advantages (b) and (c), namely Bayestraits

## Supporting Information

### S1 Table

**The organisms of this study.** A list of the 1452 organisms that we study here. For each such organism, we have a complete proteome, in the form of a fasta file.

### S1 Text

**Extracting and processing the Hogenom data.** We extract the protein sequences for all of the organisms of the Hogenom database, and process it to obtain the complete proteomes for the 1452 organisms listed in S1 Table. This procedure is detailed here in this text.

### S2 Table

**The enzymes of this study.** A list of the 2732 enzymes that we study here. The first column is the *Enzyme Commission* (EC) number, and the second is a short description of the enzyme.

### S2 Text

**Selecting the set of enzymes.** Since our study is more concerned with core metabolism than secondary metabolism, we do not consider enzymes that exclusively catalyze macromolecular reactions. How we choose such enzymes among those predictable by Priam is detailed here.

### S1 Fig

**The phylogenetic tree.** An ultrametric phylogenetic tree on the 1452 organisms listed in S1 Table.

### S3 Text

**Inferring the phylogenetic tree.** How we infer the ultrametric phylogenic tree depicted in S1 Fig.

### S2 Fig

**Expected number of gains.** On each branch of the ultrametric phylogenic tree on the 1452 organisms depicted in S1 Fig, we take the sum, over all 2732 enzymes listed in S2 Table, of the expected number of gains on the branch. The branches having the highest such values are colored red, all the way through the visible spectrum to the branches having the lowest such values, colored blue. See the legend in the upper-left corner of the figure for the correspondence of values to colors.

### S3 Fig

**Expected number of losses.** The expected number of losses are presented analgously to that of gains in S2 Fig

### S1 Script

**Modified version of Bio++.** A bash script that automatically downloads and installs a version of Bio++ modified to allow the input of probabilisitic states to several of the maximium likelihood methods that use Bio++.

### S3 Table

**Enzymes in the Kegg modules.** A list of specific enzymes, the enzymes we use in this study, that are also in Kegg modules after filtering out duplicate entries in the modules, and those modules that are of size one. The first column is the *Enzyme Commission* (EC) number, and the second is a short description of the enzyme.

### S4 Text

**Identifying the null model in the MI distribution.** Fig. 5 depicts how the mutual information (MI) between pairs of evolutionary scenarios is distributed. While some pairs of scenarios are truly concerted, with a finite sample size, some pairs will have some non-zero MI just by chance. We want to understand the nature of this later *null model*, in order to separate it from the pairs that are truly concerted. This is detailed here.

### S5 Text

**Link Communities.** We applied a link-clustering algorithm to our graph(s) *G_t_* that clusters on the *edges* of the graph, allowing communities (in terms of nodes) to overlap, because a node can appear in more than one edge cluster. The results of this are detailed here.

## Footnotes

1 More precisely, MapNH returns the expected number of gains and lossees of a character on each internal branch of a phylogeny, from which we obtain the corresponding probabilities for the respective gain and loss evolutionary scenarios.

2 Bayestraits, and therefore Discrete, is not open-source, which does not permit further investigation/improvement on these cases

